# Increasing cell size remodels the proteome and promotes senescence

**DOI:** 10.1101/2021.07.29.454227

**Authors:** Michael C. Lanz, Evgeny Zatulovskiy, Matthew P. Swaffer, Lichao Zhang, Ilayda Ilerten, Shuyuan Zhang, Dong Shin You, Georgi Marinov, Patrick McAlpine, Josh E. Elias, Jan M. Skotheim

**Author notes:** These authors contributed equally to this work (Michael C. Lanz, Evgeny Zatulovskiy).

## Abstract

Cell size is tightly controlled in healthy tissues, but it is poorly understood how variations in cell size affect cell physiology. To address this, we employed a high-accuracy mass spectrometry-based approach to measure how the proteome changes with cell size. Protein concentration changes are widespread, measurable in both asynchronous and G1-arrested cell populations, and predicted by subcellular localization, size-dependent changes in mRNA concentrations, and protein turnover. As proliferating cells grow larger, protein concentration changes typically associated with cellular senescence are increasingly pronounced. This suggests that large size is a cause rather than just a consequence of cell senescence. Consistent with this hypothesis, larger cells are prone to replicative, DNA damage-, and CDK4/6i-induced senescence. Size-dependent changes to the proteome, including those associated with senescence, are not observed when an increase in cell size is accompanied by a similar increase in ploidy. This shows that proteome composition is determined by the DNA-to-ploidy ratio rather than cell size *per se* and that polyploidization is an elegant method to generate large non-senescent cells as is commonly found in nature. Together, our findings show how cell size could impact many aspects of cell physiology through remodeling the proteome, thereby providing a rationale for cell size control and polyploidization.

## Main Text

Cells have dedicated mechanisms to control their size, which is one of the most prominent characteristics of distinct cell types ^1–3^. The link between the characteristic cell size and cell function is more obvious at the extremes of the cell size range. Red blood cells and lymphocytes need to be small to squeeze through tight spaces, while macrophages must be larger to engulf a wide range of targets. However, in the middle of the cell size range, including epithelial cells and fibroblasts, the link between cell size and function is unclear. One possibility is that these cells control their size to enhance proliferation, as their rapid turnover is a key part of their physiological function ^4,5^. Yet, even if these cells are optimized for growth and proliferation, as has been indicated ^4,6^, it is unclear why there would be an optimal cell size. As cells get larger, it has long been assumed that most proteins and RNA remain at constant concentrations ^7–14^, and organelle volumes, such as the nucleus, increase in direct proportion to cell size ^15–17^. If protein and RNA concentrations remain constant, larger cells should be capable of proportionally more biosynthesis. However, this is not the case. There is a limit to the size range of efficient biosynthesis ^18^, and excessively large cells exhibit loss of mitochondrial potential ^5^, dilution of their cytoplasm ^6^, and reduced proliferation ^19^. Moreover, recent work has demonstrated the remarkable effect even small variations in cell size can have on hematopoietic stem cell proliferation ^4^.

One possible explanation for why there is an optimal cell size for biosynthesis would be if many key cellular proteins did not remain at constant concentration as cells grow. Then, the further cells are from their target size, the more concentrations of these proteins would change, and the more growth and metabolism would deviate from the optimum. Intriguingly, investigations of the mechanisms cells use to control their size identified a class of proteins whose concentrations change with cell size. In budding yeast, human, and plant cells, key cell cycle inhibitors are not synthesized in proportion to cell size so that they are diluted by cell growth, a behavior defined as sub-scaling ^20–22^ (Fig. **1a**). Larger cells therefore have lower concentrations of cell cycle inhibitors, which promotes their division. In fission yeast, division is in part promoted by a size-dependent increase in the concentration of a cell cycle activator ^23^, a behavior defined as super-scaling ^24^ (Fig. **1a**). If this phenomenon of size-dependent protein concentration changes were widespread across the proteome it would provide an explanation for why there is an optimal cell size. This is because the further a cell would be from its target size, the further its intracellular protein concentrations would be from their optimum. To explore this possibility, we first measured how the proteome changes as a function of the natural cell size variation in asynchronously proliferating cells.

**Figure 1.**
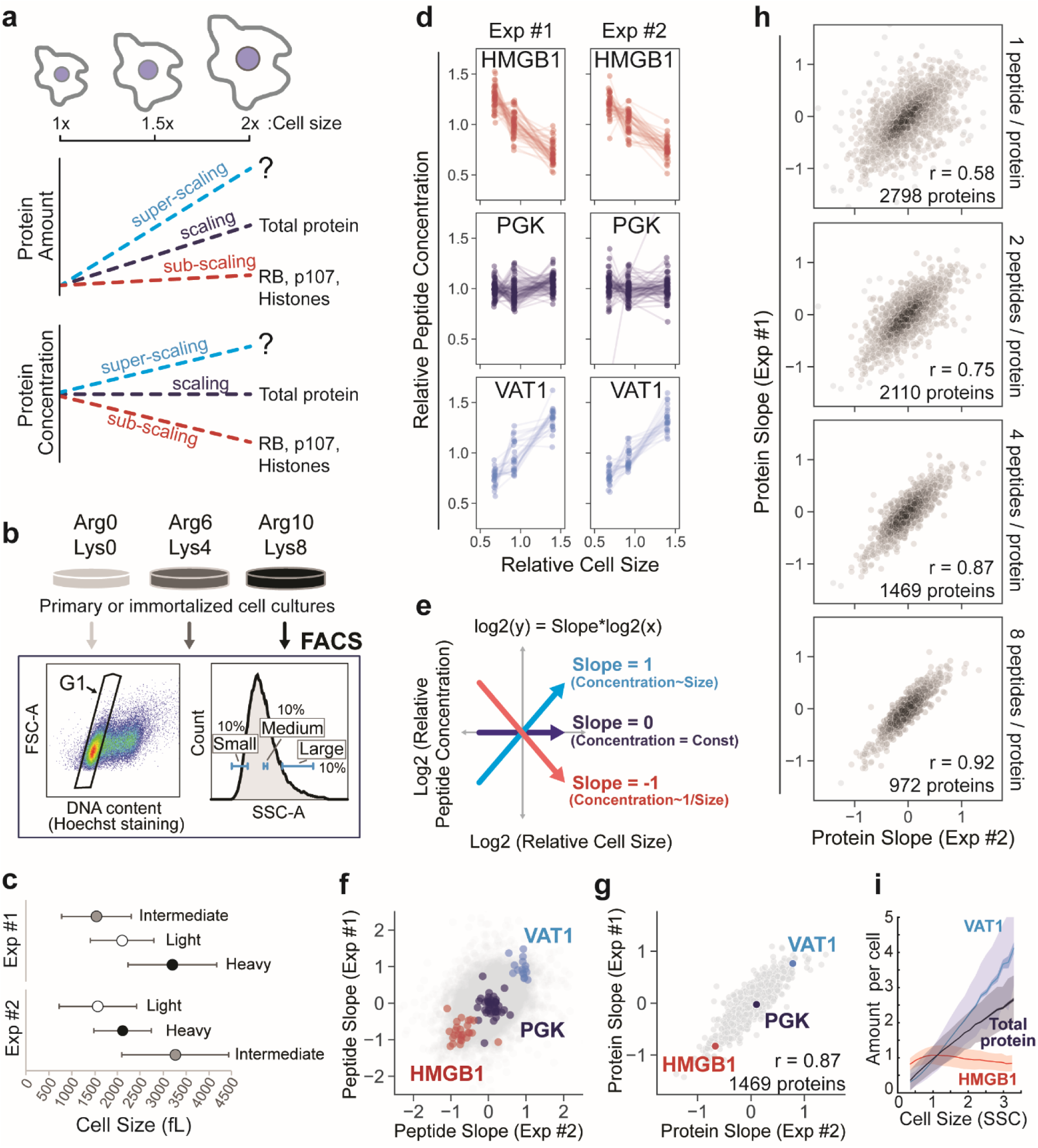
Cell size shapes the human proteome. **(a)** Schematic illustration of the potential scaling relationships between protein amount, concentration, and cell size. **(b)** Metabolically labeled HLFs cells were gated by G1 DNA content and sorted into three size bins based on the side scatter parameter (SSC) as a proxy for cell size using FACS. **(c)** The attainment of differentially sized G1 cells was confirmed using a Coulter counter. Central dots represent the mean volume for each size bin and error bars represent the standard deviation. SILAC labeling orientation was swapped for replicate experiments. **(d)** SILAC channel intensities were used to infer changes in relative peptide concentration, which are plotted for three example proteins. Each dotted line represents an independent peptide measurement. **(e)** Derivation of a slope value that describes the scaling behavior of each peptide triplet. A slope value of 1 corresponds to an increase in protein concentration that is proportional to the increase in volume and a slope of -1 corresponds to dilution (1/volume). **(f)** Peptide and **(g)** protein slope values from two replicate experiments. Only proteins with at least 4 independent peptide measurements in both experiments are shown. **(h)** Correlation of protein slope values from two replicate experiments. A threshold for the minimum number of peptide measurements per protein is indicated in each panel. **(i)** Immunofluorescence intensity measured as a function of SSC (cell size) using flow cytometry. The relative change in the total protein amount is inferred from the measurement of carboxyfluorescein succinimidyl ester (CFSE) dye. The data were binned by cell size and plotted as mean protein amounts per cell for each size bin (solid lines). Dark shaded area shows standard error of the mean for each bin, and light shaded area shows the standard deviation. A representative is shown of n=3 biological replicates for each experiment. 100,000 cells were analyzed for each sample.

To measure the scaling behavior of the human proteome (Fig. **1a**), we developed a method based on triple-SILAC (<u>S</u>table Isotope <u>L</u>abeling by <u>A</u>mino acids In <u>C</u>ell Culture) proteomics (described in Extended Data Figure 1), which enables the simultaneous measurement of thousands of individual proteins that collectively associate will all major cellular components. Asynchronously proliferating SILAC-labeled primary human lung fibroblasts (HLFs) were gated for G1 DNA content and sorted into three size bins (small, medium, and large) using fluorescence-activated cell sorting (FACS) (Fig. **1b**). SILAC labeling orientation was swapped for replicate experiments, and the attainment of differentially sized G1 cells was confirmed using a Coulter counter (Fig. **1c**, Extended Data Fig. **2a**). The SILAC channel intensities within a peptide “triplet” represent relative peptide concentrations (Extended Data Fig. **1a**), so we used the behavior of multiple independent peptide triplets to determine the size scaling relationship for individual proteins (Fig. **1d**). Rather than measure SILAC ratios, as is typically done, we calculated a slope value to describe the size scaling behavior of each peptide triplet (Fig. **1E**, Extended Data Fig. **1d-g**). Peptide triplets with positive and negative slopes represent peptides (Fig. **1f**), and therefore proteins (Fig. **1g**), whose concentrations are super- and sub-scaling, respectively, whereas triplets with a slope value near 0 maintain an approximately constant concentration as a function of G1 cell size.

We observed a continuum of size scaling behaviors across the proteome, which spanned slope values from -1 to 1, indicating that a 2-fold increase in cell size can lead to a 2-fold increase or decrease in concentration for a given protein (Fig. **1h**, Supplementary Table **1**). Several chromatin-associated High Mobility Group proteins (HMGs), including HMGB1, are diluted in larger cells and so exhibit sub-scaling behavior like what has been previously described for RB, Whi5, and histones ^21,25,26^ (Fig. **1f**). We also identified a diverse set of super-scaling proteins, such as VAT1, whose concentration increases as a function of cell size (Fig. **1d**). Setting a requirement for multiple independent measurements per protein significantly improved the correspondence between replicate experiments and yielded a high-confidence set of ∼1,500 proteins, each having at least 4 distinct peptide measurements (Fig. **1h**, Supplementary Table **1**). The size-scaling relationship was consistent across all size bins, indicating that similar concentration changes take place when cell size increases from small to medium as when it increases from medium to large (Extended Data Figure 3). Next, we validated candidate super- and sub-scaling behaviors for a subset of proteins using immunofluorescence combined with flow cytometry (Fig. **1i**, Extended Data Fig. **4**). We also confirmed that the process of cell sorting did not affect our proteome measurements (Extended Data Fig. **5a**) and that isolation of different sized cells using SSC (Side Scatter) or the total protein dye CFSE (Carboxyfluorescein succinimidyl ester) as proxies for cell size ^14,27^ yielded similar results (Extended Data Fig. **5b-d,** Supplementary Table **1**). Since larger G1 cells have, on average, spent more time in G1, it is possible that the protein concentration changes we observe reflect time in G1 rather than cell size. To control for this possibility, we selected different sized G1 cells that were synchronously released from a Thymidine-Nocodazole cell cycle arrest. We found a highly similar size-scaling relationship in our synchronous and asynchronous experiments (Extended Data Fig. **6a**, **Data S1**), indicating that cell size, not time in the G1 cell cycle phase, is driving changes to the proteome.

To examine how cell size may influence different aspects of cellular biology, we asked if different groups of related or interacting proteins exhibited similar size scaling behavior. Indeed, this was the case. For example, all 17 detected histone variants sub-scaled and had slope values near -1 (Fig. **2a**), indicating that histones are diluted by cell growth in G1 ^25,26^. Moreover, 5 of the 6 detected cathepsin proteases were strongly super-scaling and nearly doubled in concentration from the smallest to the largest cell size bin. Interestingly, we found that all individual protein subunits of the ribosome and proteasome showed small but highly consistent sub- and super-scaling behaviors respectively (Fig. **2a**) and that, more generally, proteins that are partners in a complex scale similarly (Fig. **2b**, Supplementary Table **3**). We also re-confirmed the sub-scaling behavior of the cell cycle inhibitor RB ^21^ (Extended Data Fig. **5e**, **7e**).

**Figure 2.**
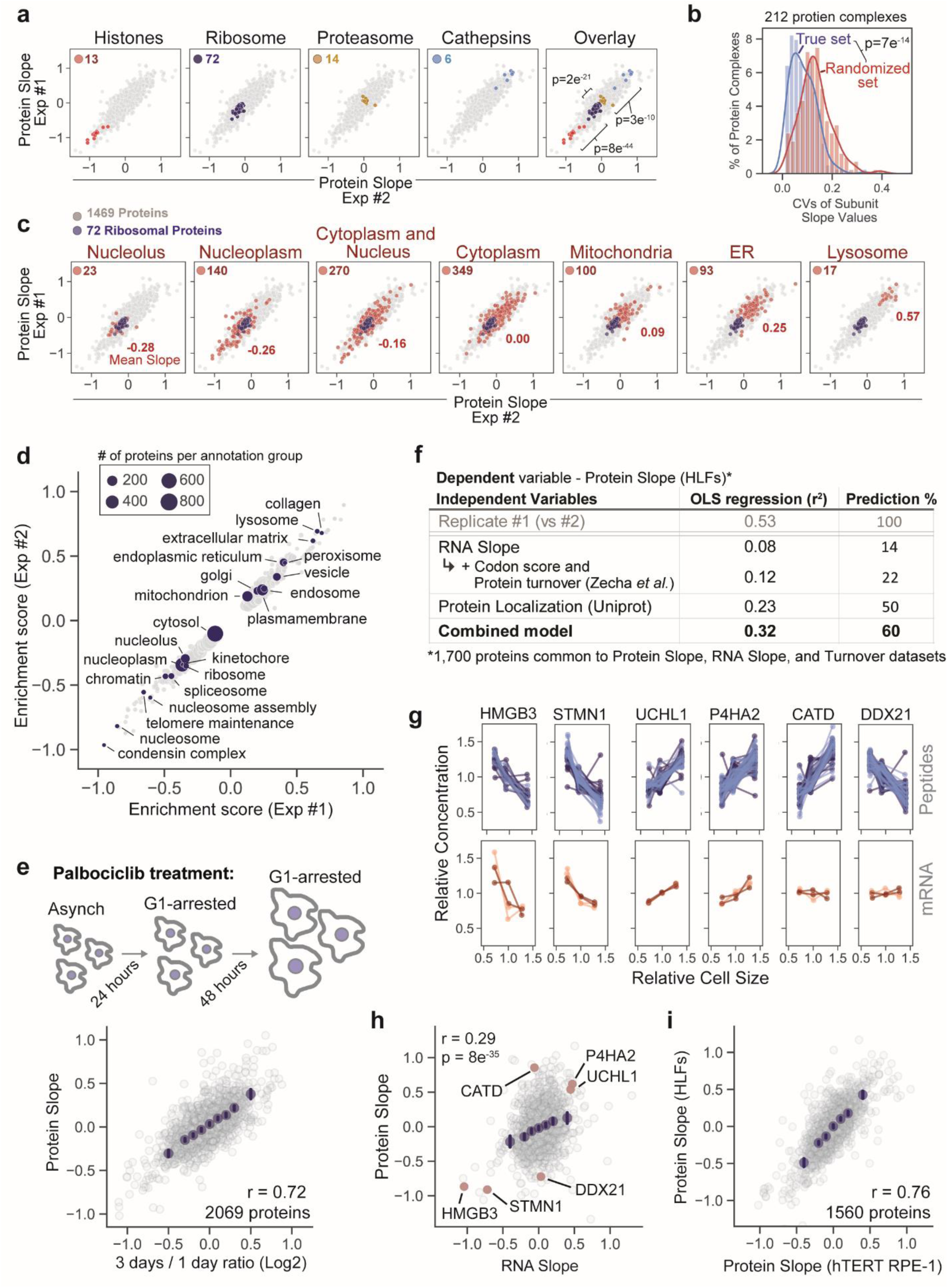
Diverse mechanisms control proteome size-dependency. **(a)** Scaling behavior of protein groups. Significance is determined by t-test between adjacent protein groups. **(b)** Variance of subunit Protein Slope values for 212 annotated protein complexes. Only proteins with cytoplasmic annotation are considered. Significance is determined by t-test between the proteins grouped by complex and a randomized control set. **(c)** Scaling behavior of proteins based on subcellular localization. The number of highlighted proteins and their average slope are indicated for each panel. Ribosomal proteins are plotted in dark blue. **(d)** 2D annotation enrichment analysis using the Protein Slope values calculated in HLF cells. Data S1 contains a complete list of enrichment scores for significantly super- or sub-scaling GO terms. **(e)** Asynchronous *hTERT RPE-1* cells were arrested in G1 with 1µM Palbociclib. Mean cells size at Day 1 and 3 was 3.5 pL and 6.5 pL, respectively. Protein ratios were determined by TMT quantitation. X-axis bins are shown in dark blue. Error bars represent the 95% confidence interval and r denotes the Pearson correlation coefficient. **(f)** Ordinary least squares regression model predicts the size scaling behavior of 1,700 individual proteins based on their subcellular localization and additional features. RNA Slope is calculated in a manner analogous to the Protein Slope using RPKM values. The benchmark for predictive accuracy (Prediction %) is determined by the correlation between biological replicate experiments. See Extended Data Fig. 9 for a full description of the model. **(g)** Size-dependent concentration changes for a representative set of proteins and their corresponding mRNA transcripts. For proteins, each connected line represents a unique peptide measurement from two biological replicate experiments (light and dark blue). For RNA, technical replicates are denoted in the same color, while biological replicates are denoted in different colors (4 replicates in total). **(h)** Correlation of size-dependent proteome and transcriptome changes. Examples in (g) are highlighted are in red. **(i)** Similarity of the cell size-dependent concentration changes between primary HLF and immortalized *hTERT RPE-1* cells. Protein Slope values for each cell type are the mean of two biological replicate experiments. Only proteins with at least 3 independent peptide measurements in each biological replicate are depicted.

We next assessed how size-scaling behavior relates to subcellular location. To do this, we first annotated the proteome based on a strict association with a single, major cellular compartment. Subcellular localization was a strong predictor of size scaling behavior, with ER- and lysosome-resident proteins becoming increasingly concentrated with size, and nucleoplasmic/nucleolar proteins becoming more dilute (Fig. **2c**, Extended Data Fig. **7a**). We observed no clear difference in scaling behavior between luminal- and membrane-associated organelle proteins (Extended Data Figure **8**). We confirmed lysosome super-scaling by measuring the lysosome-labelling dye Lysotracker and the lysosomal protein LAMP1 (Extended Data Fig. **7c**). A less stringent 2D annotation enrichment revealed a range of intermediate scaling behaviors across multiple different cellular compartments (Fig. **2d**, Supplementary Table **1**). Importantly, we confirmed that large cells generated by long-term treatment with the CDK4/6 inhibitor Palbociclib, which arrests cells in G1 but does not inhibit cell growth ^6,28^, recapitulated the proteome changes we observed in size-sorted G1 cells (Fig. **2e**, Supplementary Table **2).** In support of these findings, a proteomic analysis revealed similar size-scaling relationships when the relative protein concentration was plotted against the mean cell size for each cancer cell line in the NCI60 collection ^29^. Taken together, these analyses refute the commonly held assumption that the protein content of cellular components scales in uniform proportion to cell volume. While total protein concentration may be largely constant within a cell’s natural size range, this is not necessarily true for individual proteins.

Next, having identified size-dependent changes in concentrations of individual proteins, we sought to investigate the underlying mechanisms. We first tested whether changes in a protein’s concentration are explained by changes in the concentration of the corresponding mRNA. To do this, we performed RNA-seq on size-sorted G1 cells (Supplementary Table **4**) and calculated mRNA slope values analogous to the protein slopes described in Figure 1. We found that although there was significant correlation between the protein and RNA slopes, there was large variability in protein scaling slopes for any given RNA slope (Fig. **2f-h**). Overall, changes in mRNA concentrations explained only a minority of the variation in protein concentration size-dependency ^30^. We therefore explored additional variables using linear models. In addition to mRNA concentration, we included protein turnover ^31^, mRNA codon affinity, and subcellular localization as variables to predict size-dependent changes in protein concentration. Iterative incorporation of each parameter significantly improved the model (Fig. **2F**, Extended Data Fig. **9**). Using the correlation between biological replicates as a benchmark, we conclude that our composite model predicts ∼60% of the size-dependent variance in protein concentration and strongly supports the conclusion that the size-dependency of protein concentration is regulated both pre- and post-transcriptionally. Importantly, the size-dependency of the proteome is not specific to a single human cell type, as we found a striking degree of similarity in the proteome scaling of primary lung fibroblasts (HLF) and immortalized epithelial cells (*hTERT RPE-1*) (Fig. **2i**, Extended Data Fig. **6**).

Curiously, increasing cell size in proliferating cells is accompanied by proteomic changes normally associated with cell senescence, including the upregulation of beta-galactosidase, lysosomal proteins, and metalloproteases, and the downregulation of Ki67, HMGB proteins, and LaminB ^32^ (Fig. **3a**, Extended Data Fig. **7d**). Senescence-associated secretory phenotype (SASP) proteins ^33^ were also super scaling (Fig. **3b**). While large cell size is associated with senescence ^32^, it has generally been thought that large size results from a senescent cell’s inability to divide while at the same time maintaining cell growth. However, our experiments indicate that increasing cell size itself results in proteome changes that gradually approach those found in senescent cells and support the hypothesis that cell size *per se* may promote senescence ^4,6^ (Fig. **3c**). This is consistent with earlier reports that continued cell growth and hypertrophy are required to induce senescence in cell cycle arrested cells ^19^.

**Figure 3.**
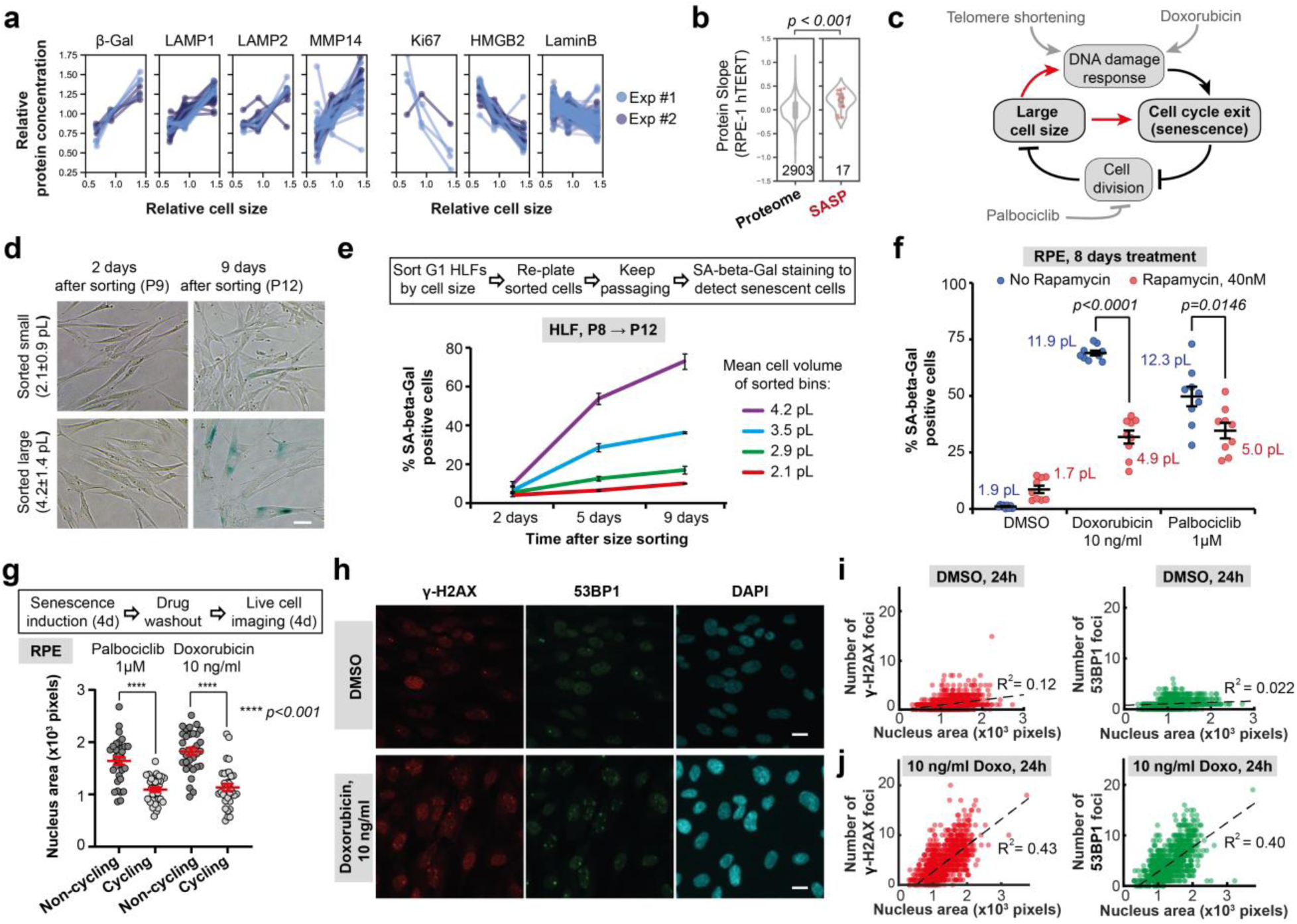
Large cell size promotes cell senescence. **(a, b)** Examples from our proteomics data set of cell size-dependent intracellular (a) and SASP (senescence-associated secretory phenotype, (b)) protein concentration changes in proliferating cells that are normally associated with senescence. **(c)** Model indicating possible relationships between DNA-related stress, large cell size, and cell senescence. **(d, e)** Replicative senescence of different-sized primary cells. Asynchronous HLFs were gated for G1 DNA content and sorted into four bins by size using FACS, then replated and stained for the senescence marker SA-beta-Gal at the indicated time points (d). Percentage of blue-stained SA-beta-Gal positive cells (e) was calculated for each time point and plotted for each sorted size bin as mean ± standard error. Cell sizes for each bin are shown as mean ± SD in (e). P denotes the cell passage number. **(f)** Effect of Rapamycin, which reduces cell growth, on the percentage (±standard error) of SA-beta-Gal positive cells in *hTERT RPE-1* cultures treated with Doxorubicin or Palbociclib for 8 days. Values shown next to each condition indicate the mean cell sizes after 8 days of treatment. **(g)** Large cell size inhibits cell cycle entry to promote senescence. *hTERT RPE-1* cells were treated with Palbociclib or a low dose of Doxorubicin for 4 days. Then, the drugs were washed out, and the cells were imaged for 4 days to identify cells that re-enter the cell cycle, and cells that remain arrested in a senescent state. Nuclear area was used as a proxy for cell size and cell cycle re-entry was determined using the fluorescent cell cycle phase reporters Cdt1-mKO2 (G1 reporter) and Geminin-mAG (S/G2/M reporter) ^41^. **(h)** Immunofluorescence staining of RPE-1 cells treated with DMSO or 10 ng/ml Doxorubicin for 24 hours against γ-H2AX (red) and 53BP1 (green), with DAPI staining shown in cyan. (**i**, **j**) Number of γ-H2AX and 53BP1 loci in RPE-1 cells treated with DMSO (i) or 10 ng/ml Doxorubicin (j) plotted against the nucleus area, which serves as a proxy for cell size. N = 33 cells for each data point in (g). SA-beta-Gal quantification for every data point in (e) included 700-1200 cells quantified from 9 different fields of view. In (i) and (j), N = 1602 and N = 1265 cells, respectively. All the experiments were performed in biological duplicates.

To test if large cell size contributes to senescence, we used primary human lung fibroblasts (HLF) that naturally senesce after 10-15 passages. We sorted HLFs into 4 cell size bins at passage #8, re-plated the cells, and then cultured them for an additional 5 passages (Extended Data Fig. **10c**). Cells that were larger at the time of sorting exhibited high levels of senescence-associated beta-galactosidase (SA-beta-Gal) staining sooner than cells that were smaller at the time of sorting (Fig. **3d,e**). This is despite the fact large and small cells were equally proliferative and did not differ in telomere length or in the mRNA concentrations of senescence markers shortly after sorting (Extended Data Fig. **10a-d**). Next, we sought to examine how cell size contributes to senescence in telomerase-immortalized human retinal pigment epithelium (*hTERT RPE-1*) cells exposed to a low dose of a DNA damaging agent (10 ng/ml Doxorubicin), which causes a stress similar to telomere shortening in primary fibroblasts. We note the proteome’s size-dependency in untreated G1 *hTERT RPE-1* cells is very similar to that of HLFs (Fig. **2i**, Extended Data Fig. **7**). Consistent with the previous result, larger cells increasingly induced the senescence marker SA-beta-Gal upon Doxorubicin treatment (Extended Data Fig. **10e,f**). Moreover, co-treatment with the mTOR inhibitor Rapamycin, which reduces cell size, significantly attenuates the SA-beta-Gal staining induced by prolonged treatment with Doxorubicin or the Cdk4/6 inhibitor Palbociclib (Fig. **3f**, Extended Data Fig. **10g, h**), a finding consistent with previous work ^34^.

So far, we have only assayed senescence using indirect markers rather than the durable cell cycle arrest that defines a senescent cell ^35^. We therefore sought to more directly test whether large cell size inhibits cell cycle re-entry following the withdrawal of two treatments previously shown to induce senescence ^6^. We treated *hTERT RPE-1* cells with Palbociclib or a low-dose of Doxorubicin for 4 days to induce senescence in a fraction of the cell population. We then washed out the drugs and imaged these enlarged cells for 4 additional days to determine which cells re-entered the cell cycle and which cells remained durably arrested (Fig. **3g**). Importantly, Palbociclib or Doxorubicin treatment arrests cells without stopping cell growth, and thus generates a population of cells with a range of large cell sizes that exhibit senescent features ^6^. Since all these cells are exposed to drug for the same duration, we can isolate cell size as a determining factor for durable cell cycle arrest. We found that the cells that were larger at the time of drug washout are more likely to remain arrested than smaller cells (Fig. **3g**). This supports the hypothesis that larger size makes cells more prone to senescence, consistent with findings for excessively large budding yeast ^6^. Large cell size may help explain the durable cell cycle arrest in response to transient exposure to a DNA damaging agent. This is because the excessive cell growth that occurs while cells delay cell cycle progression to repair DNA can inhibit cell cycle re-entry. In addition, it is possible that large cells have difficulty in replicating and repairing DNA since most DNA repair and replication factors subscale with cell size (Extended Data Fig. **11a**). To test this hypothesis, we treated the cells with a low concentration (10ng/ml) of Doxorubicin for 24 hours to induce DNA damage, and then quantified the numbers of γ-H2AX and 53BP1 loci that mark unrepaired DNA breaks in cell nuclei (Fig. **3h,i**; Extended Data Fig. **12**). We found that the number of those loci per nucleus increases with the nucleus area, supporting the idea that larger cells have more DNA damage that may promote senescence. Crucially, we are not stating that large cell size is the only driver of senescence, but rather proposing that the large size characteristic of senescent cells may itself further promote this state by inhibiting cell cycle progression and maintaining cell cycle arrest. Consistent with this hypothesis, mutations to the Rb-family proteins that reduce cell size prevent senescence of mouse embryonic fibroblasts and hematopoietic stem cells ^4,36,37^.

Our data so far are consistent with the hypothesis that large cell size inhibits cell division, and that this may be due to widespread changes in the concentrations of individual proteins as cells grow larger. The further cells are from their target size, the more protein concentrations deviate from their optimum. However, in apparent contradiction to this model, there are many examples of large animal cells that proceed through the cell cycle and maintain highly efficient cell growth ^38,39^. In many cases, such large cells are polyploid and therefore do not have an aberrant cell size-to-ploidy ratio. We therefore sought to test whether protein size-scaling is determined by cell size per se or by the cell size-to-ploidy ratio. To do this, we sought to compare the proteomes of diploid, tetraploid, and octoploid (2N, 4N, 8N) G1-phase cells (Fig. **4a,b**). To obtain G1 cells with different ploidies, we induced endoreduplication in *hTERT RPE-1* cells using a moderate dose of the Aurora kinase B inhibitor barasertib (75nM, 48 hours) ^38^ and sorted 2N, 4N, and 8N G1 cells from a mixed population using FACS (Fig. **4a**). While ploidy increases are accompanied by a corresponding increase in cell size (Fig. **4a**), proteomic analysis showed that the ploidy-sorted cells do not exhibit the same proteome changes as size-sorted cells with 2N ploidy (Fig. **4c-e**, Supplementary Table **5**). Taken together, these results indicate that the proteome’s size-dependency is largely due to changes in the cell size-to-ploidy ratio. Thus, despite being different sizes, cells of different ploidy maintain similar protein concentrations, which may be why the excessively large size of polyploid cells does not inhibit their cell cycle progression.

**Figure 4.**
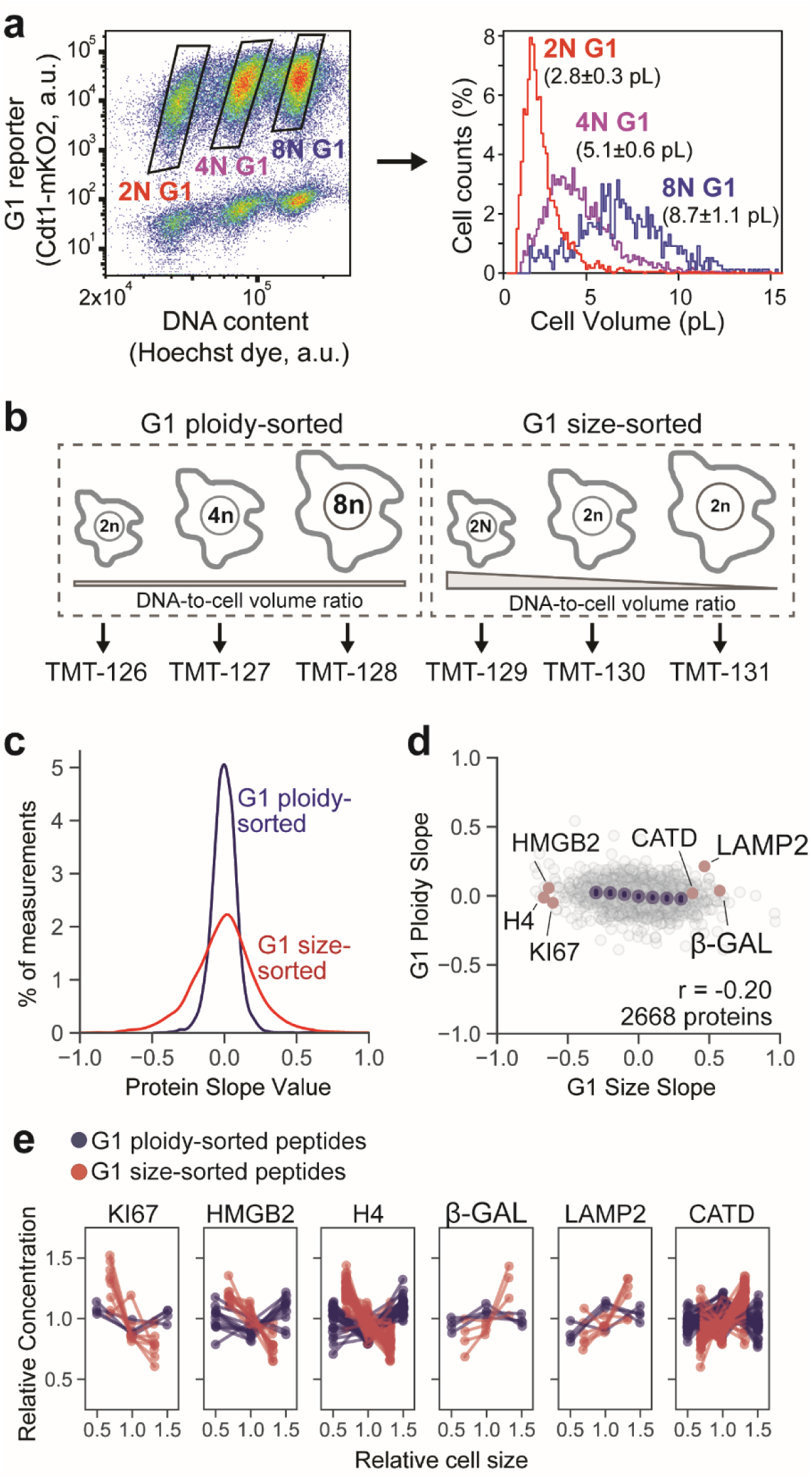
Cell volume-to-ploidy ratio drives size-dependent proteome changes. **(a)** *hTERT RPE-1* cells expressing fluorescent cell cycle reporters (Cdt1-mKO2, Geminin-mAG) were treated with an aurora kinase inhibitor barasertib (75nM, 48 hours) to partially inhibit cytokinesis. Cells were then sorted based on ploidy and G1 cell cycle phase. The attainment of differentially sized G1 cells was confirmed using a Coulter counter. The histogram shows a representative example of size distributions for sorted cells, and the numbers next to it represent mean cell size ± standard error for n=3 biological replicates. **(b)** Ploidy-sorted and Size-sorted G1 cells were isolated by FACS and their proteomes measured using TMTsixplex. Ploidy-sorted Protein Slope values were calculated by plotting the relative protein concentration against the mean cell size in the 2N, 4N, and 8N bins to obtain a slope value. **(c)** Distributions of ploidy-sorted and size-sorted Protein Slope values. Despite large increases in cell size from 2N to 8N, concentration changes were minimal. **(d)** Slightly negative correlation between ploidy-sorted and size-sorted Protein Slope values are consistent with a small increase in DNA-to-cell volume ratio in polyploid G1 cells. **(e)** DNA-to-cell volume-dependent concentration changes for a representative set of proteins. For each protein panel, dotted lines represent unique peptide measurements.

Our results here shed some light on the phenomenon of cellular senescence ^32,35,40^. While the senescent cell state has been associated with large cell size, this was mostly thought to be a passive consequence of continuing biosynthesis during cell cycle arrest. However, that such large cell sizes inhibit cell division implies the relationship between cell size and senescence can also be inverted ^4,6^. Taken together, our work suggests a model where large cell size can drive widespread changes in protein concentrations away from their optimum, which, through a yet unknown mechanism, inhibits cell division to reinforce senescence.

While it has long been thought that most protein concentrations remain constant as cells grow, this paradigm had not previously been tested using a high-throughput quantitative proteomics approach. In contradiction to the previous paradigm, many protein concentrations changed with cell size. Some proteins sub-scaled with cell size, and were diluted in larger cells, while others super-scaled with cell size so that their concentrations increased as cells grew larger. This finding is reflected in the super- and sub-scaling of the mRNA transcripts for various G1/S regulators in budding yeast ^24^. To a large extent, these diverse protein size-scaling behaviors could be predicted from a linear model based on mRNA concentration, protein half-life, and subcellular localization, which indicates the importance of both transcriptional and post-transcriptional size-scaling mechanisms.

Our observation here that most protein concentrations change as cells grow provides a rationale for why many cells control their size. If the proteome content that supports optimal cell growth is only found near the target cell size, then the further a cell deviates from its target size, the further protein concentrations will be from their growth-supporting optimum. While a small change in the concentration of a single protein may not significantly affect cell physiology, the cumulative effect of thousands of small protein concentration changes could account for the drastic drop in the efficiency of biosynthesis in large cells ^4,6,19^. Moreover, since proteome concentrations mostly follow the cell size-to-ploidy ratio rather than cell size *per se*, polyploidization is an elegant mechanism for organisms to generate large cells capable of efficient protein synthesis as is commonly found in nature.

## Acknowledgements

We thank Shicong Xie for assistance with python-related data analysis and Marcus Smolka for use of his HPLC for HILIC pre-fractionation. We thank Robert Brooks, Chuck Sherr, Christine Jacobs-Wagner, Marcus Smolka, Jette Lengefeld, and members of the Skotheim lab for helpful feedback on the manuscript. We also thank Frank McCarty and Stanford SUMS for assistance in mass spec related data acquisition. Cell sorting was performed on an instrument in the Shared FACS Facility purchased by Parker Institute for Cancer Immunotherapy.

## Author contributions

EZ conceived and coordinated the project. MCL and EZ designed the study. EZ performed and analyzed all cell biology and flow cytometry experiments, prepared and sorted cells for proteomics. MCL processed all proteomics samples. MCL, LZ, and JEE conceived data acquisition strategy. LZ and PM acquired mass spectrometry data. MCL and İİ analyzed proteomics data. MCL and SZ derived the linear models. EZ, MPS and GM analyzed the RNAseq data. DY analyzed lysosome scaling with cell size. JMS supervised the study. MCL, EZ and JMS interpreted the data and wrote the manuscript.

## Supplementary Materials

### Materials and Methods

#### Cell Culture

Recently isolated primary fetal human lung fibroblasts (HLF) were purchased from Cell Applications, telomerase-immortalized retinal pigment epithelium (*hTERT RPE-1*, here also referred to as RPE-1 for brevity) cells were obtained from the Stearns laboratory at Stanford. All cells were cultured at 37°C with 5% CO_2_ in Dulbecco’s modification of Eagle’s medium (DMEM) with L-glutamine, 4.5 g/l glucose and sodium pyruvate (Corning), supplemented with 10% heat-inactivated fetal bovine serum (FBS, Corning) and 1% penicillin/streptomycin.

#### Fluorescence-activated cell sorting (FACS)

Fluorescence-activated cell sorting was used to sort live cells by their size and cell cycle phase. To do this, the cells were harvested from dishes by trypsinization, stained with 20 µM Hoechst 33342 DNA dye in PBS for 15 minutes at 37°C, and then sorted on a BD FACSAria Fusion flow cytometer. Consecutive SSC-A over FSC-A, and FSC-H over FSC-A gates were used to isolate single cells. Then, G1 cells were gated by DNA content (Hoechst staining). Finally, we collected the 10% smallest and 10% largest cells, as well as another 10% of the cells near the average size using the gating based on SSC-A signal. During sorting, all cell samples and collection tubes were kept at 4°C. To determine the cell size distributions of the collected samples, aliquots were taken from each sorted size bin and measured on a Z2 Coulter counter (Beckman). Sorted cells were used for mRNA or protein isolation, or re-plated for assessing senescence dynamics.

#### Stable Isotope Labeling In Cell culture (SILAC)

Cells for SILAC were cultured in special Lysine/Arginine-free DMEM for SILAC (Thermo Scientific) with 10% dialyzed heat-inactivated FBS (HyClone) and penicillin/streptomycin. These cultures were supplemented with “light”, “intermediate”, or “heavy” versions of Lysine (0.8mM) and Arginine (0.4mM) (Cambridge Isotope Laboratories) ^42^. The “light” (Agr0 Lys0) version of the media contained L-Arginine and L-Lysine built with normal ^12^C and ^14^N isotopes; the “intermediate” (Arg6 Lys4) version had L-Arginine containing six ^13^C atoms and L-Lysine containing four deuterium atoms; the “heavy” (Arg10 Lys8) version had L-Arginine containing six ^13^C and four ^15^N atoms and L-Lysine containing six ^13^C and two ^15^N atoms. Proline (200 mg/l) was added to the media to prevent conversion of isotope-coded Arginine into Proline in cells. We confirmed that cell proliferation is not impaired in our SILAC medium (Extended Data **2b**). To ensure complete labelling, the cells were cultured in SILAC media for 5 passages (approximately 10 doublings) prior to the experiments.

#### LC-MS/MS sample preparation - SILAC

See Table S3 for a complete list of proteomic experiments. Small, medium and large cells sorted by FACS were pelleted by centrifugation at 500xg for 10 minutes and lysed for 40 minutes on ice in RIPA lysis buffer (Abcam) containing a protease and phosphatase inhibitor cocktail. SILAC-labeled different sized cells were mixed prior to lysis in order to minimize handling error in protein extraction, proteolytic digestion, and peptide desalting. Cell lysates were cleared by centrifugation at 15000xg for 30 minutes at 4℃. The mixed lysates were then denatured in 1% SDS, reduced with 10mM DTT, alkylated with 5mM iodoacetamide, and then precipitated with three volumes of a solution containing 50% acetone and 50% ethanol. Proteins were re-solubilized in 2 M urea, 50 mM Tris-HCl, pH 8.0, and 150 mM NaCl, and then digested with TPCK-treated trypsin (50:1) overnight at 37°C. Trifluoroacetic acid and formic acid were added to the digested peptides for a final concentration of 0.2%. Peptides were desalted with a Sep-Pak 50mg C18 column (Waters). The C18 column was conditioned with 5 column volumes of 80% acetonitrile and 0.1% acetic acid and washed with 5 column volumes of 0.1% trifluoroacetic acid. After samples were loaded, the column was washed with 5 column volumes of 0.1% acetic acid followed by elution with 4 column volumes of 80% acetonitrile and 0.1% acetic acid. The elution was dried in a Concentrator at 45°C.

#### LC-MS/MS sample preparation - TMT

Lysis, denaturation, reduction, and precipitation for SILAC analysis was the same for TMT analysis (working solution of Iodoacetamide was dissolved in HEPES rather than Tris buffer). Our method for TMT labeling was adapted from Zecha et al. ^43^ and the Thermo TMT10plex™ Isobaric Label Reagent Set Protocol. In brief, acetone precipitated samples were resuspended in 100µm TEAB and digested O/N with TPCK trypsin (50:1) in the absence of Tris or Urea. After digestion, peptide concentration was ∼1µg/ul in 100µM TEAB for all samples. 20µg of peptide was labeled using 100µg of Thermo TMT10plex™ in a reaction volume of 25µl for 1 hour. The labeling reaction was quenched with 8µL of 5% hydroxylamine for 15 minutes. Labeled peptides were pooled, acidified to a pH of ∼2 using drops of 10% formic acid, and desalted with a Sep-Pak 50mg C18 column as described above.

#### HILIC fractionation - SILAC

Desalted peptide samples were reconstituted in 80% acetonitrile and 1% formic acid and fractionated by hydrophilic interaction liquid chromatography (HILIC) with a TSK gel Amide-80 column (2 mm x 150 mm, 5 µm; Tosoh Bioscience). 90 second fractions were collected between 10 and 25 minutes of the gradient. Three solvents were used for the gradient: buffer A (90% acetonitrile), buffer B (80% acetonitrile and 0.005% trifluoroacetic acid), and buffer C (0.025% trifluoroacetic acid). The gradient used consists of a 100% buffer A at time = 0 min; 88% of buffer B and 12% of buffer C at time = 5 min; 60% of buffer B and 40% of buffer C at time = 30 min; and 5% of buffer B and 95 % of buffer C from time = 35 to 45 min in a flow of 150 µl/min. HILIC fractions were dried in a SpeedVac and reconstituted in 0.1% trifluoroacetic acid. A total of 10 fractions were collected and pooled back into 5 fractions (1-6, 2-7, 3-8, 4-9, 5-10).

#### High-pH reverse phase fractionation - TMT

TMT-labeled peptides (Experiment from Figure S5D) were fractionated using a Pierce™ High pH Reversed-Phase Peptide Fractionation Kit. The eight default fractions were pooled back to 4 fractions (1-5, 2-6, 3-7, 4-8). Dried peptides were reconstituted in 0.1% formic acid.

#### LC-MS/MS data acquisition - SILAC

Peptide samples were analyzed using a Fusion Lumos mass spectrometer (Thermo Fisher Scientific, San Jose, CA) equipped with Dionex Ultimate 3000 LC systems (Thermo Fisher Scientific, San Jose, CA). Peptides were separated by capillary reverse phase chromatography on a 25 cm reversed phase column (100 µm inner diameter, packed in-house with ReproSil-Pur C18-AQ 3.0 m resin (Dr. Maisch GmbH)). Liquid chromatography was performed using a two-step linear gradient with 4–25 % buffer B (0.1% (v/v) formic acid in acetonitrile) for 90 min followed by 25-40 % buffer B for 10 min. Data was acquired in top 20 data dependent mode. Full MS scans were acquired in the Orbitrap mass analyzer with a resolution of 120,000 (FWHM) and m/z scan range of 340-1500. Selected precursor ions were subjected to fragmentation using higher-energy collisional dissociation (HCD) with quadrupole isolation, isolation window of 1.6 m/z, and normalized collision energy of 30%. HCD fragments were analyzed in the Orbitrap mass analyzer with a resolution of 15,000 (FWHM). Fragmented ions were dynamically excluded from further selection for a period of 15 seconds. The AGC target was set to 400,000 and 50,000 for full FTMS scans and FTMS2 scans, respectively. The maximum injection time was set to 50 ms for full FTMS scans and dynamic for FTMS2 scans.

#### LC-MS/MS data acquisition - TMT

Desalted TMT-labeled peptides were analyzed on a Fusion Lumos mass spectrometer (Thermo Fisher Scientific, San Jose, CA) equipped with a Thermo EASY-nLC 1200 LC system (Thermo Fisher Scientific, San Jose, CA). Peptides were separated by capillary reverse phase chromatography on a 25 cm column (75 µm inner diameter, packed with 1.6 µm C18 resin, AUR2-25075C18A, Ionopticks, Victoria Australia). Electrospray Ionization voltage was set to 1550 volts. Peptides resulting from on-bead digestion were resuspended in 10 µL of 0.1% formic acid. 2 µL was introduced into the Fusion Lumos mass spectrometer using a two-step linear gradient with 6– 33% buffer B (0.1% (v/v) formic acid in 80% acetonitrile) for 145 min followed by 33-45% buffer B for 15 min at a flow rate of 300 nL/min. Column temperature was maintained at 40°C throughout the procedure. Xcalibur software (Thermo Fisher Scientific) was used for the data acquisition and the instrument was operated in data-dependent mode. Survey scans were acquired in the Orbitrap mass analyzer over the range of 380 to 1800 m/z with a mass resolution of 70,000 (at m/z 200). Ions were selected for fragmentation from the 10 most abundant ions with a charge state of either 2, 3 or 4 and within an isolation window of 2.0 m/z. Selected ions were fragmented by Higher-energy Collisional Dissociation (HCD) with normalized collision energies of 27 and the tandem mass spectra was acquired in the Orbitrap mass analyzer with a mass resolution of 17,500 (at m/z 200). Repeated sequencing of peptides was kept to a minimum by dynamic exclusion of the sequenced peptides for 30 seconds. For MS/MS, the AGC target was set to 1e5 and max injection time was set to 120ms. Relative changes in peptide concentration were determined at the MS3- level by isolating and fragmenting the 4 most dominant MS2 ion peaks.

#### Spectral searches - TMT and SILAC

All raw files were searched using the Andromeda engine ^44^ embedded in MaxQuant (v1.6.7.0) ^45^. See Table S4 for a complete summary of the search parameters used for the SILAC- and TMT-labeled peptide fragments. In brief, 3 label SILAC search was conducted using Maxquant’s default Arg6/10 and Lys4/8. For TMT searches, a Reporter ion MS3 search was conducted using 10plex TMT isobaric labels. For both TMT and SILAC searches, variable modifications included oxidation (M) and protein N-terminal acetylation. Carbamidomthyl (C) was a fixed modification. The number of modifications per peptide was capped at five. Digestion was set to tryptic (proline-blocked). Peptides were “Re-quantified”, and maxquant’s match-between-runs feature was not enabled. Database search was conducted using the Uniprot proteome - Human_UP000005640_9606. Minimum peptide length was 7 amino acids. FDR was determined using a reverse decoy proteome ^46^.

#### Peptide quantitation - SILAC

Our SILAC analysis pipeline uses the peptide feature information in MaxQuant’s “evidence.txt” output file. Each row of the “evidence.txt” file represents an independent peptide triplet measurement. Contaminant and decoy peptide identifications were discarded. Peptides without signal in any of the three SILAC channels were also excluded. Peptide triplets (each row in the “evidence.txt” table) are assigned to a protein based on MaxQuant’s “Leading razor protein” designation. For each peptide triplet, the fraction of ion intensity in each SILAC channel was calculated by dividing the “Intensity L/M/H” column by the “Intensity” column. SILAC channels were normalized by adjusting the fraction of ion intensity in each channel by the median for all measured peptides (see Extended Data **1b,c**). After normalization, the relative signal difference between the SILAC channels for each peptide triplet was plotted against the normalized cell size for each of the bins of isolated G1 cells.

For each peptide, we calculated its slope as follows (mean squared error filtering):

Y_1,2,3_ = Relative signal in each SILAC channel (order based on labeling orientation)

Avg. size = (mean volume of small bin + mean volume of medium bin + mean volume of large bin) / 3

x_1_ = (Mean volume of small size bin) / Avg. size

x_2_ = (Mean volume of medium size bin) / Avg. size

x_3_ = (Mean volume of large size bin) / Avg. size

Based on the expectation that our experimental conditions would not result in large, non-linear changes in protein expression, we exclude peptide triplets whose three data points did not loosely fit a linear regression line. Linear regressions on the ∼50,000 triplets/experiment were performed using np.polyfit in Python. Regressions with a mean squared error > 0.075 were excluded. Because this step significantly improved the overall data quality (Extended Data **1f,g**), we concluded that our filtering method mostly excludes peptide triplets contaminated by analytical interference or that are near the noise floor.

Individual peptide measurements were consolidated into a protein level measurement using python’s groupby.median. Peptides with the same amino acid sequence that were identified as different charge states or in different fractions were considered independent measurements. We summarize the size scaling behavior of individual proteins as a slope value derived from a regression (similar to what is described above for individual peptides), and each protein slope value is based on the behavior of all detected peptides.

For a given protein, we calculate its cell size-dependent slope as follows:

y_i_ = Relative signal in the i^th^ SILAC channel (median of all corresponding peptides in this channel)

x_i_ = same normalized cell size x_i_as for the peptide slope calculations above

The protein slope value was determined from a linear fit to the log-transformed data using the equation:

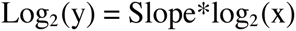

Variables were log-transformed so that a slope of 1 corresponds to an increase in protein concentration that is proportional to the increase in volume and a slope of -1 corresponds to 1/volume dilution. Pearson r and p values for correlation analyses were calculated using scipy’s pearsonr module in python.

#### Protein annotations

Protein annotations in Figure 2 were sourced from Uniprot columns named “Subcellular location [CC]” or “Protein names” ^47^. For Figure 2c, protein localization was strictly parsed so that each annotated protein belongs to only one of the designated groups. Proteins with 2 or more of the depicted annotations were ignored (except for the “Cytoplasm / Nucleus” category, which required a nuclear and cytoplasmic annotation). 2D annotation enrichment in Fig. 2d was performed using Perseus ^48^.

#### Peptide quantitation - TMT

Our TMT analysis pipeline uses the peptide feature information in MaxQuant’s “evidence.txt” output file. Each row of the “evidence.txt” file represents an independent MS^3^ TMT measurement. Contaminant and decoy peptide identifications were discarded. Peptides without signal in any of the TMT channels were also excluded. TMT peptide measurements were assigned to protein based on MaxQuant’s “Leading razor protein” designation. For each peptide triplet, the fraction of ion intensity in each TMT channel was calculated by dividing the “Reporter ion intensity” column by the sum of all reporter ion intensities. TMT channels were normalized by adjusting the fraction of ion intensity in each channel by the median for all measured peptides (similar to the SILAC normalization in Extended Data **1b,c**). After normalization, the relative signal difference between the TMT channels for each peptide triplet was plotted against the normalized cell size for each of the bins of isolated G1 cells. Slope values in Figure S5 were derived in a manner analogous to the Slope values calculated in the SILAC experiments. Pearson r and p values for correlation analyses were calculated using SciPy’s pearsonr module in python.

#### Protein complex analysis

Protein complex annotations were derived from the CORUM database (“Core Complexes” with 3512 entries). We calculated the Coefficient of Variation (CV) between the Protein Slope values of subunits in annotated protein complexes. To do so, all slope values were normalized to a value between 0 and 1. Slope values were then grouped for each protein complex. Only proteins with a minimum of 5 independent peptide measurements (between two replicate experiments) and protein complexes with at least three quantified subunits were considered in the analysis. To generate a randomized dataset, quantified proteins (*i.e.*, proteins with Protein Slope values) within each protein complex were replaced with randomly selected proteins using the random function from the NumPy library. CV is calculated for both real and randomized datasets and compared with ttest_ind (T-test) from scipy.stats library. Only cytoplasmic proteins and protein complexes were considered in order to avoid biases from the differential scaling observed in different subcellular compartments.

#### OLS linear regression model

Multiple linear regression analysis was performed using the statsmodels module in python. The prediction of size scaling behavior was based on the 1,700 proteins shared between the protein turnover (HeLa cells) ^31^, RNA Slope, and Protein Slope datasets (at least 2 peptides / protein). Independent variables for codon affinity, RNA Slope, and Protein turnover (T50%) were each independently standardized by subtracting all values by the dataset’s mean and then dividing by the dataset’s standard deviation. The codon affinity refers to the binding affinity at the 3rd codon gene is the average of the low affinity codon percentage within each amino acid, weighted by the percentage of the amino acid in that gene. The subcellular localization variable was based on Uniprot’s “Subcellular location [CC]” annotations and entered as a binary value for each compartment (1 if a protein possessed an annotation and 0 if it did not). Only subcellular compartments that provided nonredundant predictive power were ultimately included in the model. A constant value was added to the regression equation using the add_constant function in statsmodels. We set the benchmark for predictive accuracy (Prediction %) as the correlation between biological replicate experiments (Protein Slope from Exp #1 vs Exp #2). See Figure S9 for more details on the model and its coefficients.

#### RNA extraction and sequencing

To compare the transcriptomes of different-sized G1 cells, primary HLFs were arrested in the G1 phase of the cell cycle by a 24-hour treatment with 1μM of the Cdk4/6 inhibitor Palbociclib and then sorted into size bins using FACS. To extract RNA, each sample of sorted HLF cells was split into two technical replicates each of which contained 200,000 HLF cells, which were then mixed with 100,000 *D. melanogaster* S2 cells as a spike-in. Each sample was then pelleted and RNA was extracted using Direct-zol™ RNA Microprep Kit (Zymo Research). mRNA was enriched using the NEBNext Poly(A) mRNA Magnetic Isolation Module (NEB, #E7490). The NEBNext Ultra II RNA Library Prep Kit for Illumina^®^ (NEB, #E7775) was then used to prepare libraries for paired-end (2×150 bp) Illumina sequencing (Novogene). Two independent biological replicates of each sample were collected and for each biological replicate, two technical replicates (*i.e.*, separate lysis, library prep, and sequencing) were processed. Approximately 40 million reads were sequenced per replicate.

#### RNAseq data processing

RNA samples contained a mixture of *H. sapiens* and *D. melanogaster* spike-in. A combined *H. sapiens* and *D. melanogaster* genome file was created using the hg38 and dm6 versions of the respective genomes and a combined transcriptome annotation was created using the *H. sapiens* gene models from the v29 version of the GENCODE annotation ^50^ and the BDGP6 *D. melanogaster* gene models from ENSEMBL release 90 ^51^. For the purposes of RNA-seq data quality evaluation, genome browser track generation, and calculating the hg38-to-dm6 ratio, reads were aligned against the combined genomes and combined annotated set of splice junctions using the STAR aligner (version 2.5.3a; settings: --limitSjdbInsertNsj 10000000 -- outFilterMultimapNmax 50 --outFilterMismatchNmax 999 –outFilterMismatchNoverReadLmax 0.04 --alignIntronMin 10 --alignIntronMax 1000000 --alignMatesGapMax 1000000 -- alignSJoverhangMin 8 --alignSJDBoverhangMin 1 --sjdbScore 1 --twopassMode Basic -- twopass1readsN -1) ^52^. Read mapping statistics and genome browser tracks were generated using custom Python scripts. For quantification purposes, reads were aligned as 2×50mers in transcriptome space against an index generated from the combined annotations described above using Bowtie (version 1.0.1; settings: -e 200 -a -X 1000) ^53^. Alignments were then quantified using eXpress (version 1.5.1) ^54^ before effective read count values and TPM (Transcripts Per Million transcripts) were then separated for each genome and renormalized TPMs were calculated with respect to only the *H. sapiens* transcripts.

#### Flow cytometry

For flow cytometry analysis, cells were grown on dishes to ∼50% confluence and harvested by trypsinization. The cells were then fixed with 3% formaldehyde for 10 minutes at 37°C, and then permeabilized with 90% methanol for 30 minutes on ice. Fixed and permeabilized cells were washed once with PBS, blocked with 3% BSA in PBS for 30 minutes at 37°C, and then stained with primary antibodies for 2 hours at 37°C. We used the following primary antibodies: HMGB1 (Abcam, ab79823), HMGN2 (CST, 9437S), RPLP0 (Sigma, SAB1402899), Actin (Sigma-Aldrich, A2103), UCHL1 (CST, 13179S), VAT1 (Santa Cruz, sc-515705), LAMP1 (CST, 9091), alpha-Tubulin (Abcam, ab6160). After the primary antibodies, the cells were washed twice with a wash buffer (1% BSA in PBS + 0.05% Tween® 20), stained with the fluorophore-conjugated secondary antibodies Alexa Fluor 488 goat anti-mouse (Life Technologies A11029), Alexa Fluor 594 goat anti-rabbit (Life Technologies A11037), and Alexa Fluor 405 goat anti-rat (Abcam ab175673) at 1:1000 dilution for 1 hour at 37°C, and then washed twice again. After this treatment, the cells were resuspended in PBS containing 3 µM DAPI for DNA staining, incubated for 10 minutes at room temperature, and then analysed on a Attune NxT flow cytometer (Thermo Fisher). To compensate for the nonspecific background staining with antibodies, we measured the fluorescence of cells stained with nonspecific Isotype Control antibodies. We then performed a linear regression of this nonspecific background signal with the cell size, and subtracted the background fluorescence corresponding to the cell’s size from the actual fluorescence signal measured for each cell. For total protein staining, live cells were resuspended in PBS, then the CellTrace CFSE dye (CarboxyFluorescein Succinimidyl Ester, Thermo Fisher) was added at 5 μM concentration, incubated for 30 minutes at 37°C. The dye was then washed out with FBS-containing DMEM, and the cells were pelleted and resuspended in PBS for the flow cytometry or for the fixation and antibody staining. For the Lysotracker staining, the cells were harvested and re-suspended in growth media at a concentration of 10^6^ cells/mL. Then, Lysotracker Red DND-99 (Thermo Fisher) was added at a concentration of 75nM and incubated at 37°C for 30 min. Cells were spun down and re-suspended for analysis or additional staining. For plotting the flow cytometry data, all protein amounts and cell size values were normalized to their means. To characterize the degree of size-dependency of protein amounts, we fit a line to the flow cytometry data after normalizing these data to mean values. We performed at least three biological replicates for each experiment that measured 100,000 cells.

To compare the proliferation efficiency of different-sized cells shortly after FACS sorting, the cells were left to settle on culture dishes for 2 days, then cultured in the presence of 1µM EdU for 24 hours to label all the cells that underwent replication within this time period. Cells were then stained using a Click-iT™ EdU Alexa Fluor™ 647 Flow Cytometry Assay Kit (Molecular Probes), following the manufacturer’s protocol, and analyzed by flow cytometry. To compare the telomere lengths of different-sized cells, passage #8 HLF cells were stained using a Flow-FISH Telomere PNA Kit (Agilent DAKO) according to the manufacturer’s protocol and analyzed by flow cytometry.

#### Senescence-associated beta-galactosidase (SA-beta-Gal) staining

To detect senescent cells, the RPE-1 or HLF cells on a dish were stained using the Senescence β-Galactosidase Staining Kit (Cell Signaling Technology) following the manufacturer’s protocol. Briefly, live cells on a dish were washed once with PBS and fixed with 1x Fixation solution for 10 minutes at room temperature, then rinsed twice with PBS, and stained with β-Galactosidase Staining Solution for 48 hours at 37°C in a dry incubator (no CO_2_). The cells were then imaged on an EVOS M5000 imaging system to obtain a colored brightfield image. The obtained images were quantified manually, by a blinded investigator, to determine the percentage of senescent cells, *i.e.*, the cells that have a pronounced blue staining.

#### Live cell fluorescence microscopy

In preparation for imaging, cells were seeded on 35-mm glass-bottom dishes (MatTek) at low density and incubated overnight at 37°C and 5% CO_2_. Then, the cells were moved to a Zeiss Axio Observer Z1 microscope equipped with an incubation chamber and imaged for 96 hours ^55^. Brightfield and fluorescence images were collected from three dishes at multiple positions every 20 minutes using an automated stage controlled by the Micro-Manager software. We used a Zyla 5.5 sCMOS camera, which has a large field of view allowing us to track motile cells within a field of view for long durations, and an A-plan 10x/0.25NA Ph1 objective. To distinguish G0/G1 and S/G2/M cells in time lapse imaging experiments, we used RPE-1 cells expressing the fluorescent cell cycle reporters mKO2-hCdt1 (G1), and mAG-hGeminin (S/G2) ^41^. These reporters were introduced into RPE-1 cells using a lentivirus vector, and the positive, fluorescent population of cells was isolated using fluorescence-activated cell sorting.

#### Immunofluorescent staining for microscopy

Cells were seeded on a 35-mm collagen-coated glass-bottom dish (MatTek) one day before starting the cell treatments. Cells were treated with 10 or 100ng/ml Doxorubicin or with DMSO for 24 hours, and then stained with antibodies. For the staining, cells were fixed with 4% formaldehyde for 10 minutes at room temperature, permeabilized with 0.2% Triton™ X-100 (Sigma-Aldrich) for 15 minutes at 4°C, and then blocked with 3% BSA in PBS. Then, the cells were incubated with primary rabbit anti-53BP1 antibodies (Novus Biologicals, NB100304, 1:2000 dilution) and mouse anti-γ-H2AX antibodies (Sigma Millipore, 05-636-I, 1:200 dilution) overnight at 4°C, washed twice with PBS, and then incubated with conjugated Alexa Fluor 647 goat-anti-mouse (Invitrogen, A32728) and Alexa Fluor 488 goat-anti-rabbit (Invitrogen, A32731) secondary antibodies at 1:1000 for 1 hour at room temperature. The cells were washed twice with PBS, incubated with 300 nM DAPI for 5 minutes at room temperature, and then imaged using a Zeiss Axio Observer Z1 microscope with an A-plan 10x/0.25NA objective. γ-H2AX and 53BP1 loci were quantified from microscopy images using standard tools implemented in FIJI software.

---

**Extended Data Figure 1.**
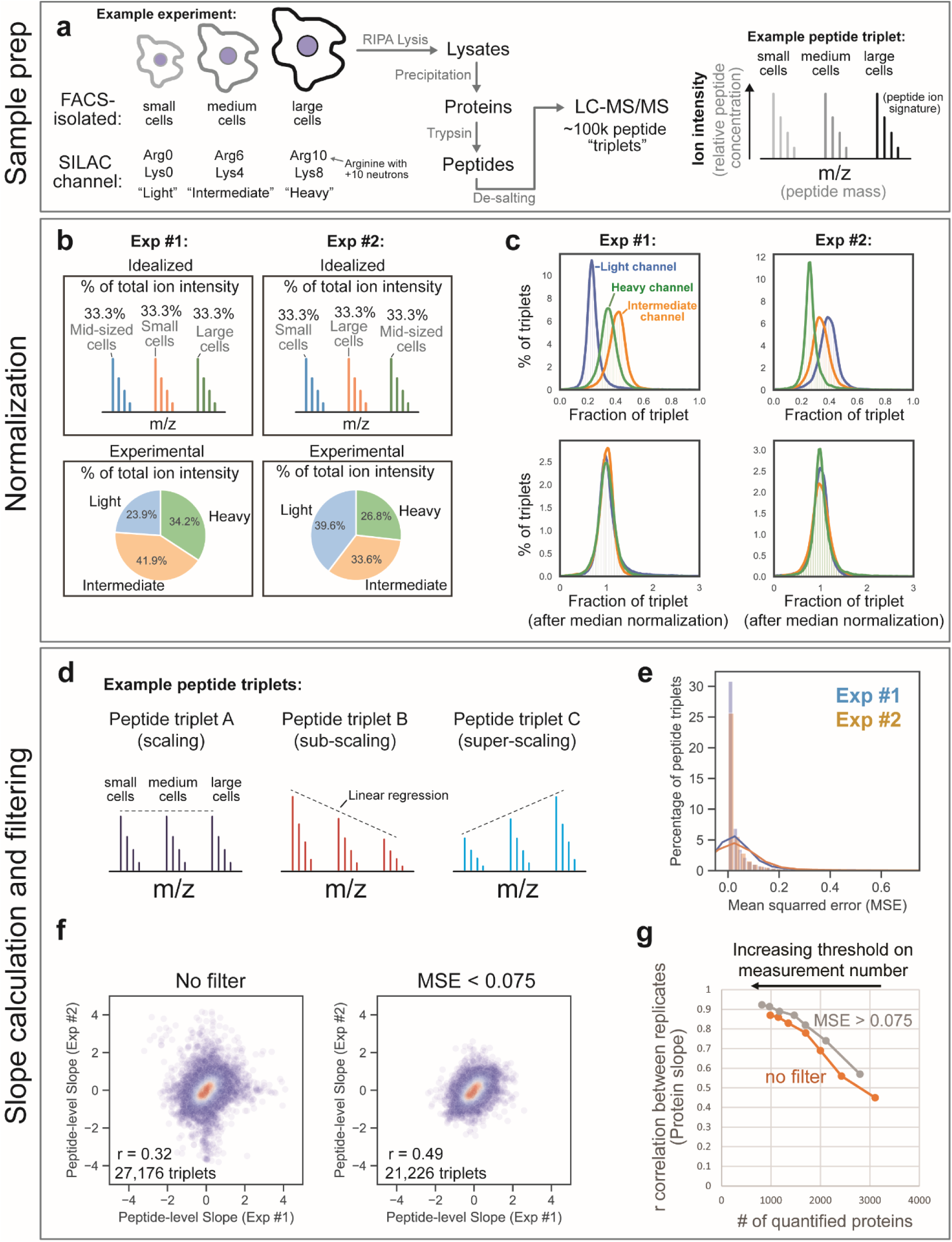
A SILAC proteomics method to measure how the proteome changes as a function of G1 cell size. **(a)** Human cells were metabolically labeled in cell culture, sorted by G1 cell size using FACS, and subjected to proteomic analysis. **(b)** Differences in the amount of total protein contributed from the small-, medium-, or large-cell size populations were normalized using the signal proportionality from the light, intermediate, and heavy channel of each peptide triplet. Small, medium, and large cells were mixed prior to lysis, so the amount of protein in each SILAC channel is uneven. Rather than normalize L/H and L/M SILAC ratios separately, we normalize all three channels together so that the values in our dataset represent relative changes to each peptide. **(c)** For each individual peptide triplet, we determined the fraction of the triplet’s total ion intensity present in each SILAC channel. The distributions of these fractions were then adjusted by the median (see methods for a complete description of the normalization process). **(d)** Peptide slope values are calculated from a linear regression of the relative ion intensity in each SILAC channel and mean cell size. Mean cell size was determined by Coulter counter prior to mixing and lysis. **(e)** Distribution of mean squared error values for peptide triplet regressions (∼50,000 per experiment). The mean squared error was used to track the linear fit of each peptide regression. **(f)** Correlation of peptide slopes calculated from biological replicate experiments before and after applying a filter for mean squared error (MSE). 27,176 unique peptide measurements were identified in both replicate experiments. A unique peptide measurement is defined by the peptide sequence, modification state, charge state, and the fraction number (the fraction number is only considered if the experiment was pre-fractioned and multiple fractions were analyzed). **(g)** Filtering peptides by mean squared error from a linear fit improves data quality. MSE filtering improves the correlation of protein slope values derived from biological replicates, and the improvement is consistent across different thresholds of measurement confidence (*i.e.*, peptide measurements per protein).

**Extended Data Figure 2.**
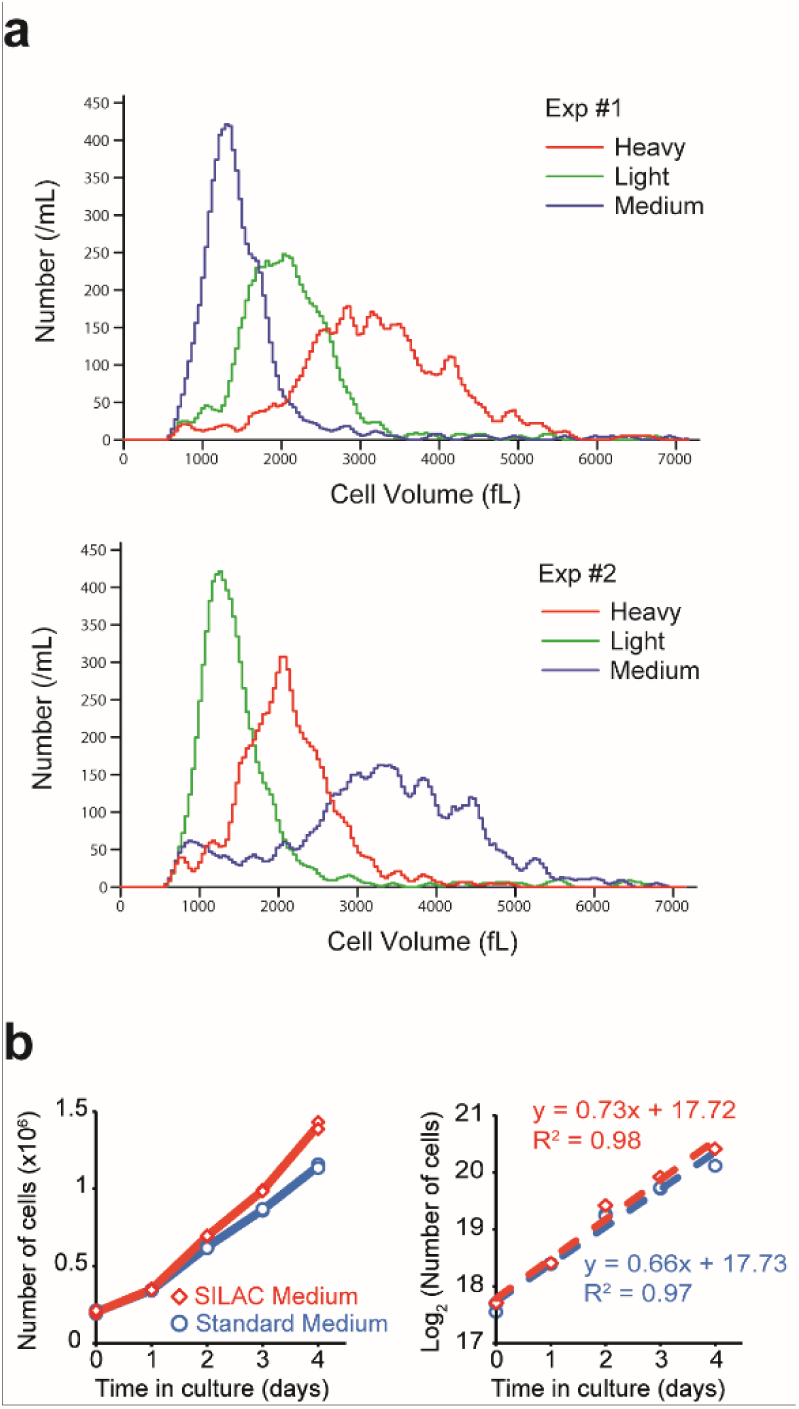
Supporting information for SILAC proteomics data shown in Figure 1: **(a)** Coulter counter measurement of HLF cells isolated by FACS. Small, medium, and large cell populations are colored based on the SILAC labeling orientation for the two replicate experiments in Figure 1. See Table S3 for cell size measurements for all proteomic experiments. **(b)** HFL primary cell proliferation rates in SILAC vs standard medium.

**Extended Data Figure 3.**
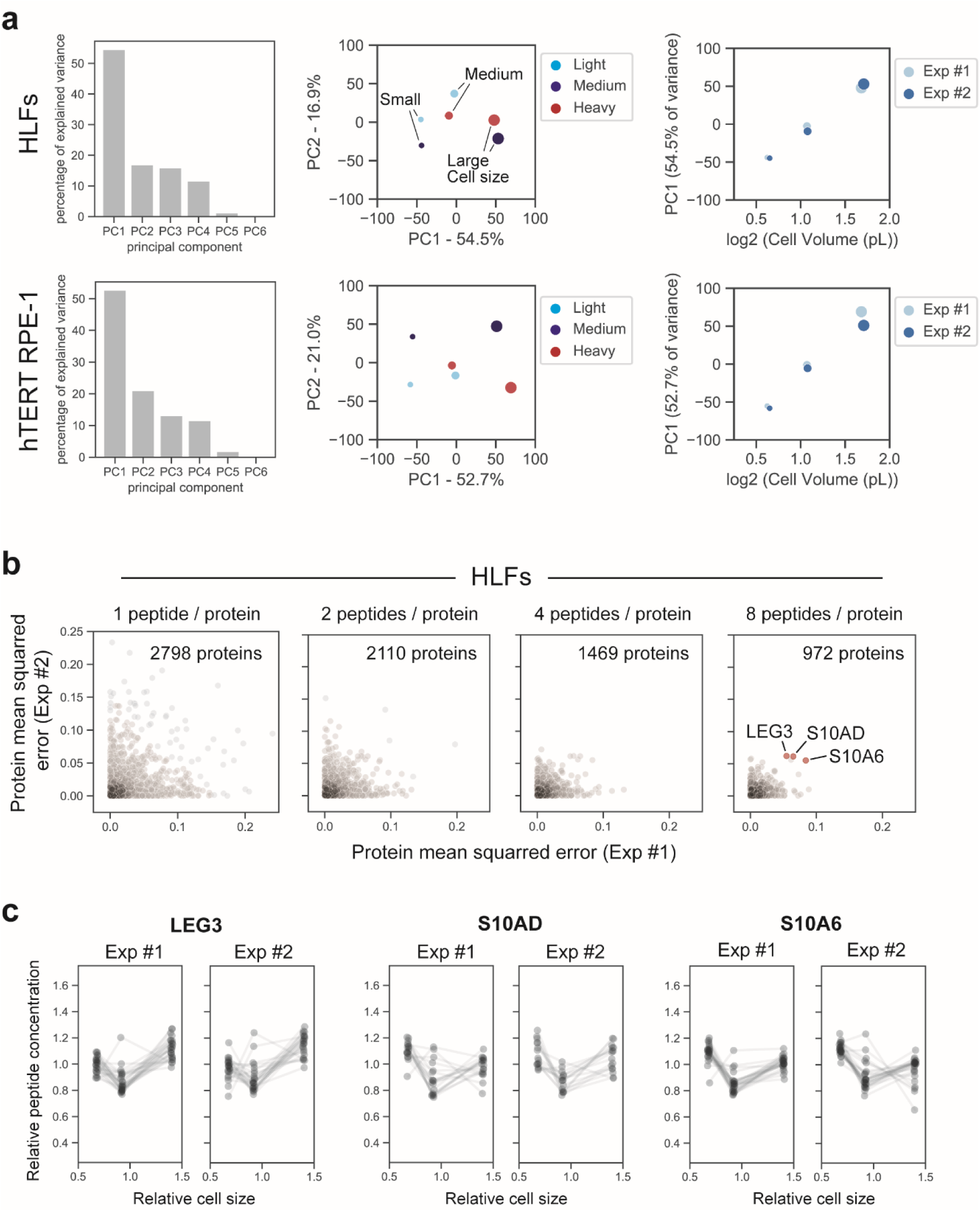
Cell size-dependent changes to concentrations in the proteome are mostly linear: **(a)** Principal component analysis (PCA) of the replicate experiments performed using HLF and hTERT RPE-1 cell lines. Data frame input for the PCA contained the relative SILAC ion channel intensity (“light”, “medium”, and “heavy”) for every measured protein in each experiment (after filtering for MSE). Dot size represents the mean cell size corresponding to each SILAC channel. PC1 represents the majority of variance in both experiments and correlates with the change in cell size. **(b)** Correlation for the mean squared error (MSE) of the Protein Slope regression from two biological replicates. A threshold for the minimum number of peptide measurements per protein is increased from left to right. Because very few proteins have reproducible large MSE values, we conclude that most proteins scale linearly with G1 cell size. **(c)** Peptide-level measurements for the few proteins with non-linear scaling are plotted below.

**Extended Data Figure 4.**
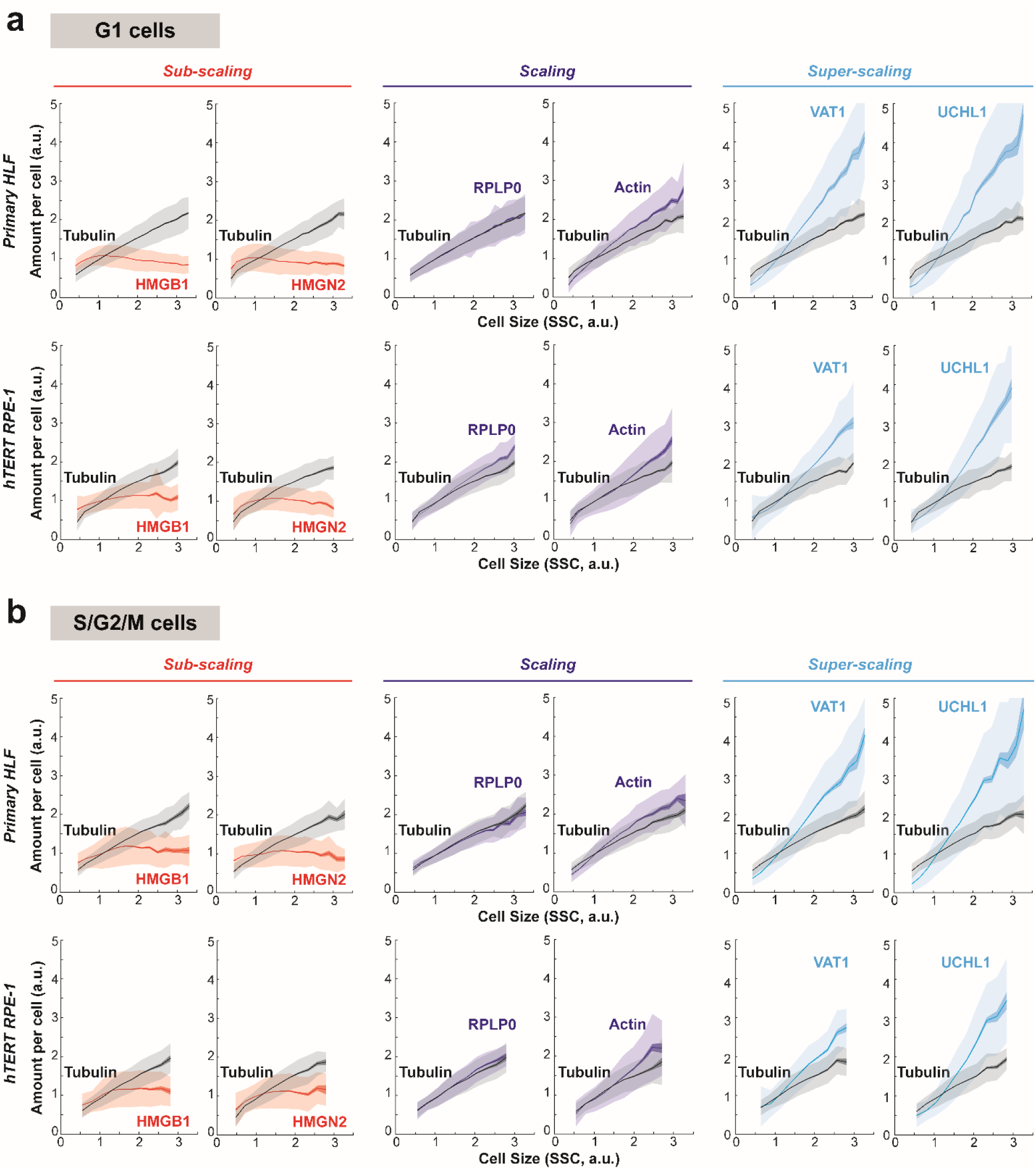
Validation of protein size-scaling behaviors using flow cytometry: Cycling HLF and RPE-1 cells were fixed and stained with antibodies against subscaling (HMGB1, HMGN2), scaling (RPLP0, beta-Actin), and superscaling proteins (VAT1, UCHL1). Alpha-tubulin is used as an internal control for each sample. Using flow cytometry, G1 and non-G1 cells were gated by DNA content (DAPI dye) and analyzed separately in panels **(a)** and **(b)**, respectively. The data were binned by cell size (SSC, the side scatter parameter) and plotted as mean protein amounts per cell for each size bin (solid lines). Dark shaded area shows standard error of the mean for each bin, and light shaded area shows the standard deviation. A representative is shown of n=3 biological replicates for each experiment. 100,000 cells were analyzed for each sample.

**Extended Data Figure 5.**
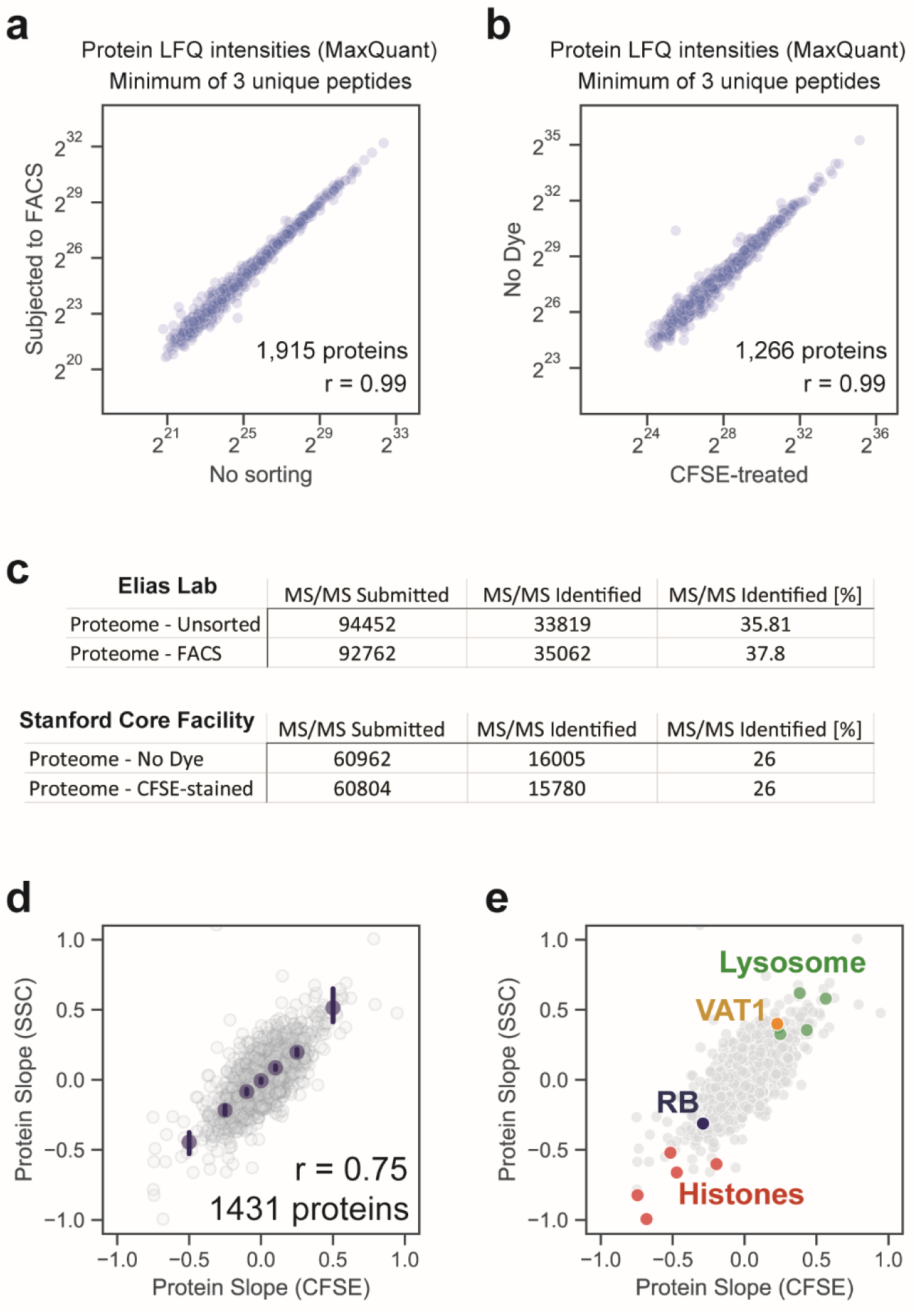
Controls indicating that cell sorting does not affect proteomics measurements: **(a)** A dish of HLF cells were split into two equal parts, and one half was run through the FACS machine while the other half sat on ice. MaxQuant LFQ was used to determine whether cells subjected to FACS exhibited altered proteomes. The strong correlation between the proteomes of sorted and unsorted cells suggests that FACS did not appreciably bias our measurements. **(b)** MaxQuant LFQ was used to compare proteome samples from CFSE-stained and unlabeled cells. The strong correlation between the proteomes of CFSE-treated and untreated cells suggests that using a total protein dye does not appreciably bias our measurement. **(c)** Peptide discovery is not impacted by FACS or CFSE staining. **(d)** RPE-1 cells with G1 DNA content were sorted using total protein / cell (CFSE stain) or side scatter to achieve 3 bins of different sized cells. Protein Slope values derived from cells sorted by total protein and side scatter are compared. A select set of proteins from the comparison in (d) are highlighted in **(e)**.

**Extended Data Figure 6.**
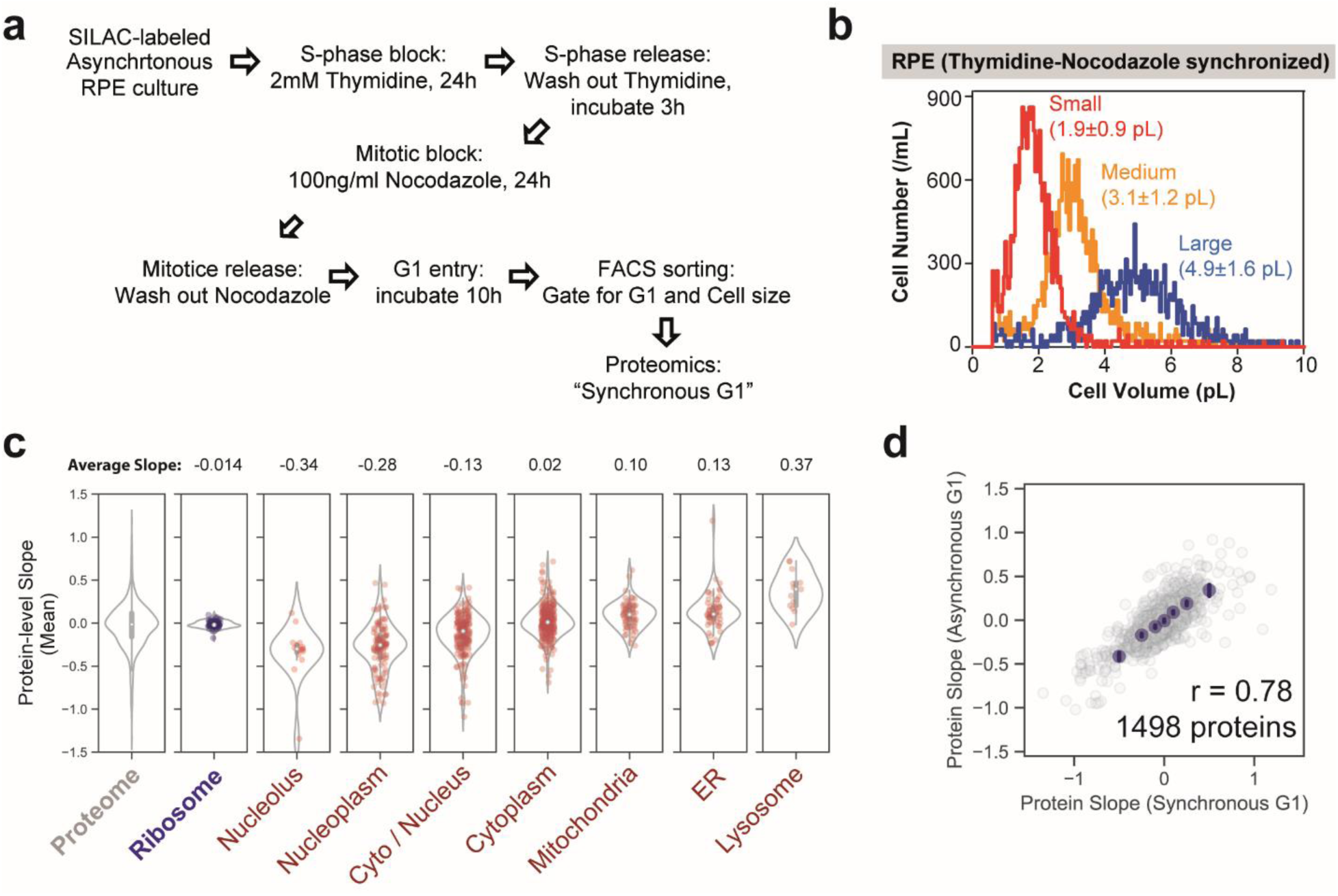
Changes to the proteome are primarily driven by cell size and not cell age: **(a)** Metabolically labeled *hTERT RPE-1* cells were synchronously released into the cell cycle following a Thymidine-Nocodazole cell cycle arrest. **(b)** A similar distribution of G1 sizes were isolated from cells synchronously released into G1 and from asynchronous cultures. **(c)** Protein slopes were calculated as described in Figure 1. Only proteins with at least 4 peptide measurements in both replicate experiments are considered for the violin plots. **(d)** Correlation of Protein Slope values calculated from RPE-1 cells synchronously released into G1 and from asynchronous cultures.

**Extended Data Figure 7.**
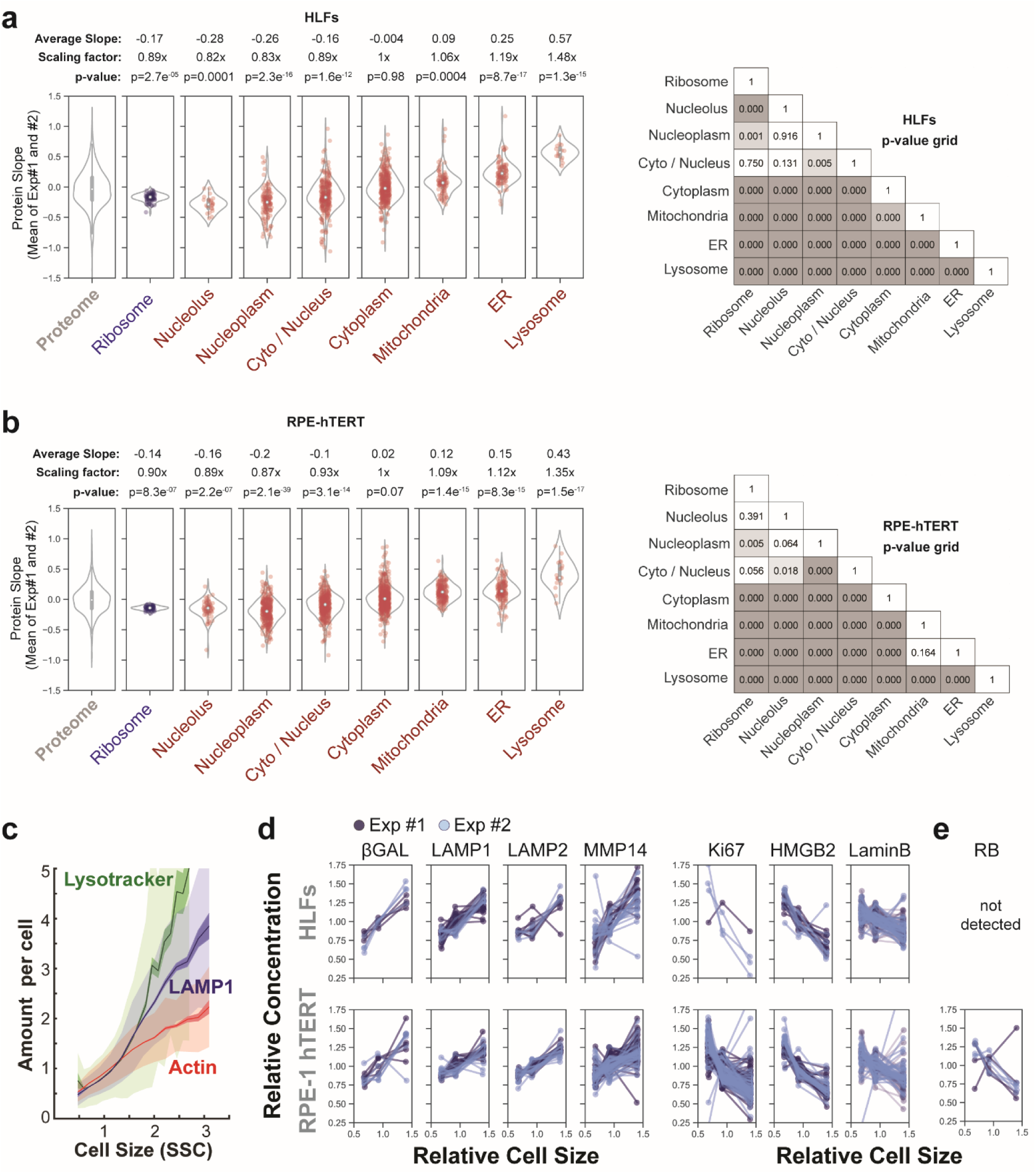
Size scaling of proteome content in an epithelial (RPE-1 hTERT) and a fibroblast (HLF) cell line: **(a)** Distribution of slopes derived from HLF cells for proteins associated with the indicated compartments. Violin plots depict the average slopes for the proteins highlighted in Figure 2b. P-values above the violin plots are derived from a t-test between the indicated protein group and the rest of the dataset. t-tests comparing the slopes for each group of proteins are visualized in a grid format. **(b)** Replicate experiment using the immortalized *hTERT RPE-1* cell line was performed as in (a). **(c)** Validation of lysosome super-scaling with cell size using flow cytometry. Both the lysosomal protein LAMP1 and the Lysotracker dye amount increase with cell size faster than Actin, which is a proxy for total protein. The data for G1 *hTERT RPE-1* cells were binned by cell size (SSC, the side scatter parameter) and plotted as mean protein amounts per cell for each size bin (solid lines). Dark shaded area shows standard error of the mean for each bin, and light shaded area shows the standard deviation. A representative is shown of n=5 biological replicates for each experiment. About 100,000 cells were analyzed for each sample. **(d)** Examples from our proteomics data set of cell-size-dependent protein concentration changes in proliferating cells that are normally associated with senescence. **(e)** RB is diluted with increasing G1 cell size.

**Extended Data Figure 8.**
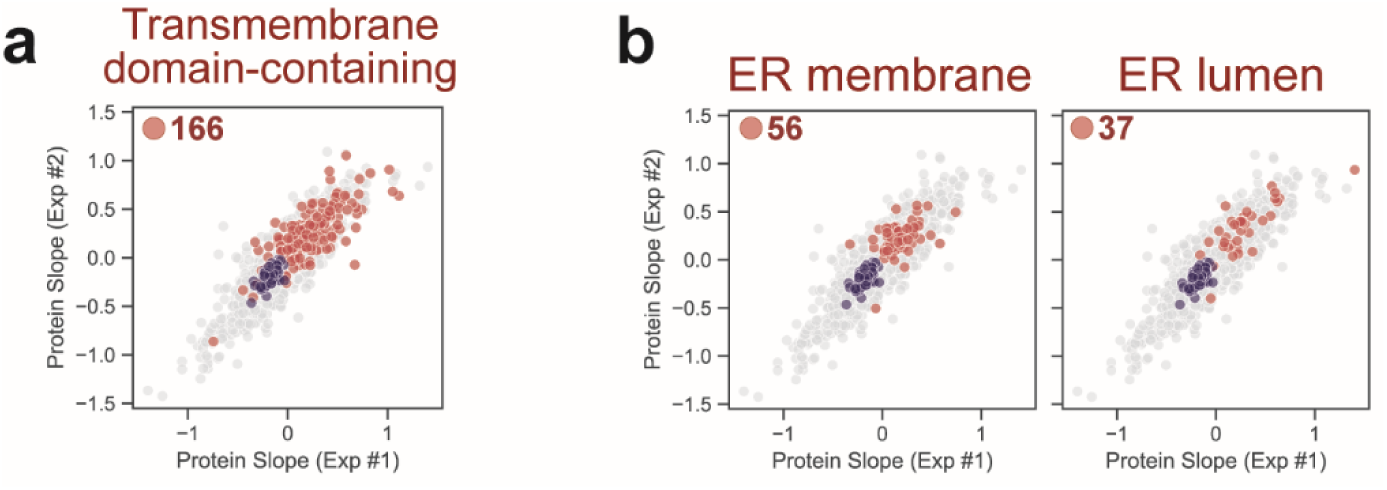
Scaling behavior of lumenal and membrane-associated proteins: **(a)** Scaling behavior of proteins predicted to contain a transmembrane domain (Uniprot’s “Transmembrane” annotation column). **(b)** Scaling behavior of ER proteins that are annotated to be either membrane-associated or not (*i.e.*, lumenal).

**Extended Data Figure 9.**
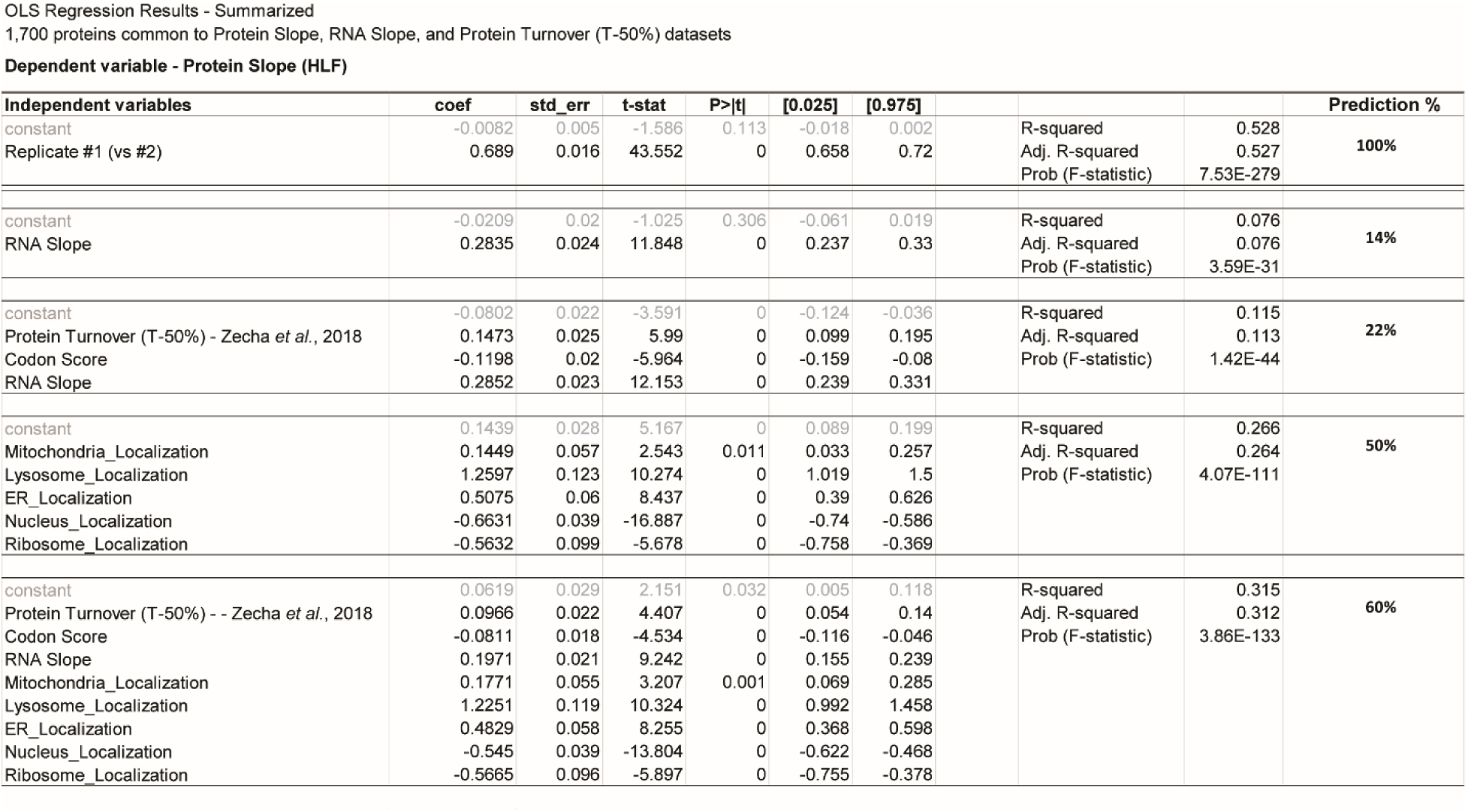
Linear regression analysis predicts size scaling behavior of individual proteins: The prediction of size scaling behavior was based on the 1,700 proteins that are in the published protein turnover dataset (HeLa cells) ^31^, as well as our RNA Slope, and Protein Slope datasets (at least 2 peptides / protein) that we report here. Independent variables for codon affinity, RNA Slope, and Protein turnover (time to replace 50% of a given protein species) were each independently standardized by subtracting all values by the dataset’s mean and then dividing by the dataset’s standard deviation. The subcellular localization variable was based on Uniprot’s “Subcellular location [CC]” annotations and entered as a binary value for each compartment (1 if a protein possessed an annotation and 0 if it did not). Only subcellular compartments that provided nonredundant predictive power were ultimately included in the model. A constant value was added to the regression equation using the add_constant function in statsmodels. We set the benchmark for predictive accuracy (Prediction %) as the correlation between biological replicate experiments, *i.e.*, Protein Slope from Exp #1 vs Exp #2.

**Extended Data Figure 10.**
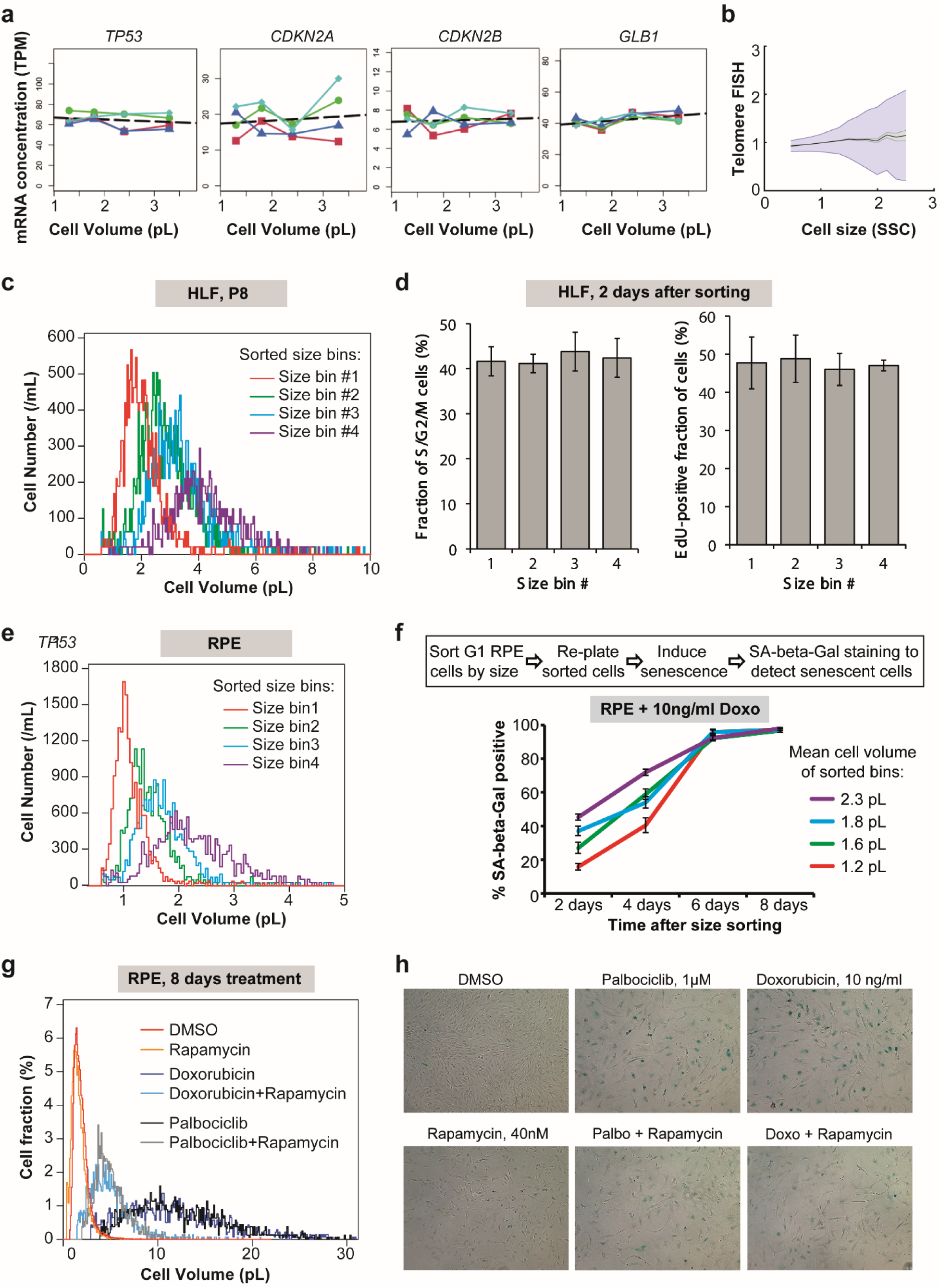
Analysis of senescence markers in different-sized cells: **(a)** Transcript levels of key senescence reporter genes in size-sorted HLF cells. G1 HLF cells were sorted into four size bins using FACS. The concentrations of *TP53, CDKN2A, CDKN2B,* and *GLB1* mRNAs were then determined by RNAseq and plotted against the mean cell size for each bin. Each colored line represents one of four replicates. **(b)** Flow-FISH Telomere PNA staining in passage #8 HLF cells plotted against cell size. **(c)** Cell size distributions for the four bins of FACS-sorted HLF cells that were then re-plated and passaged to determine replicative senescence dynamics (see Fig. 3d,e), as measured with a Coulter counter. **(d)** Percentage of S/G2/M phase cells (left) and percentage of EdU-positive cells after 24 hours labeling with 1µM EdU in HLF cells, 2 days after sorting G1 cells by cell size. **(e-f)** Asynchronous *hTERT RPE-1* cells were gated for G1 DNA content and sorted into four bins by size using FACS (e). Then the sorted cells were replated and cultured in the presence of the DNA damaging agent Doxorubicin (10 ng/ml), and then stained for SA-beta-Gal at the indicated time points to determine the DNA-damage-induced senescence dynamics (f). **(g-h)** Cell size distributions (g) and SA-beta-Gal staining images (h) of the RPE-1 cells treated for 8 days with DMSO, Palbociclib or Doxorubicin, in the presence or absence of Rapamycin to determine the effect of cell size reduction on senescence dynamics (see Fig. 3f). Each of the experiments in (b-h) was performed in two biological replicates.

**Extended Data Figure 11.**
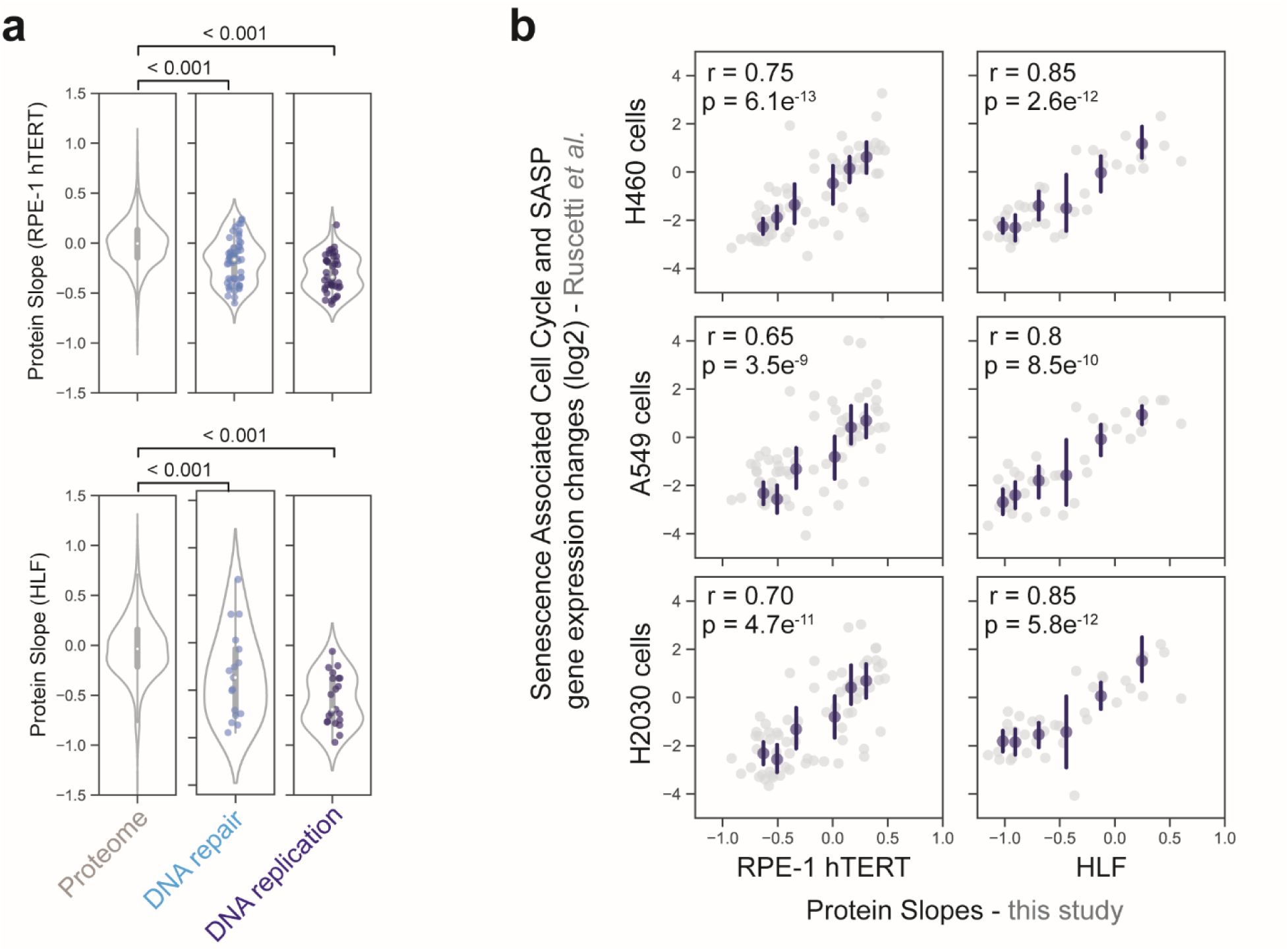
Classes of sub- and super-scaling proteins in HLF and hTERT RPE-1 cells: **(a)** Slopes for proteins grouped by Uniprot GO annotations for DNA repair (GO:0006281) and DNA replication (GO:0006260). **(b)** Size-dependent proteome changes from this study correlate with the senescence-associated SASP and cell cycle gene expression changes defined by Ruscetti, *et al.* ^33^.

**Extended Data Figure 12.**
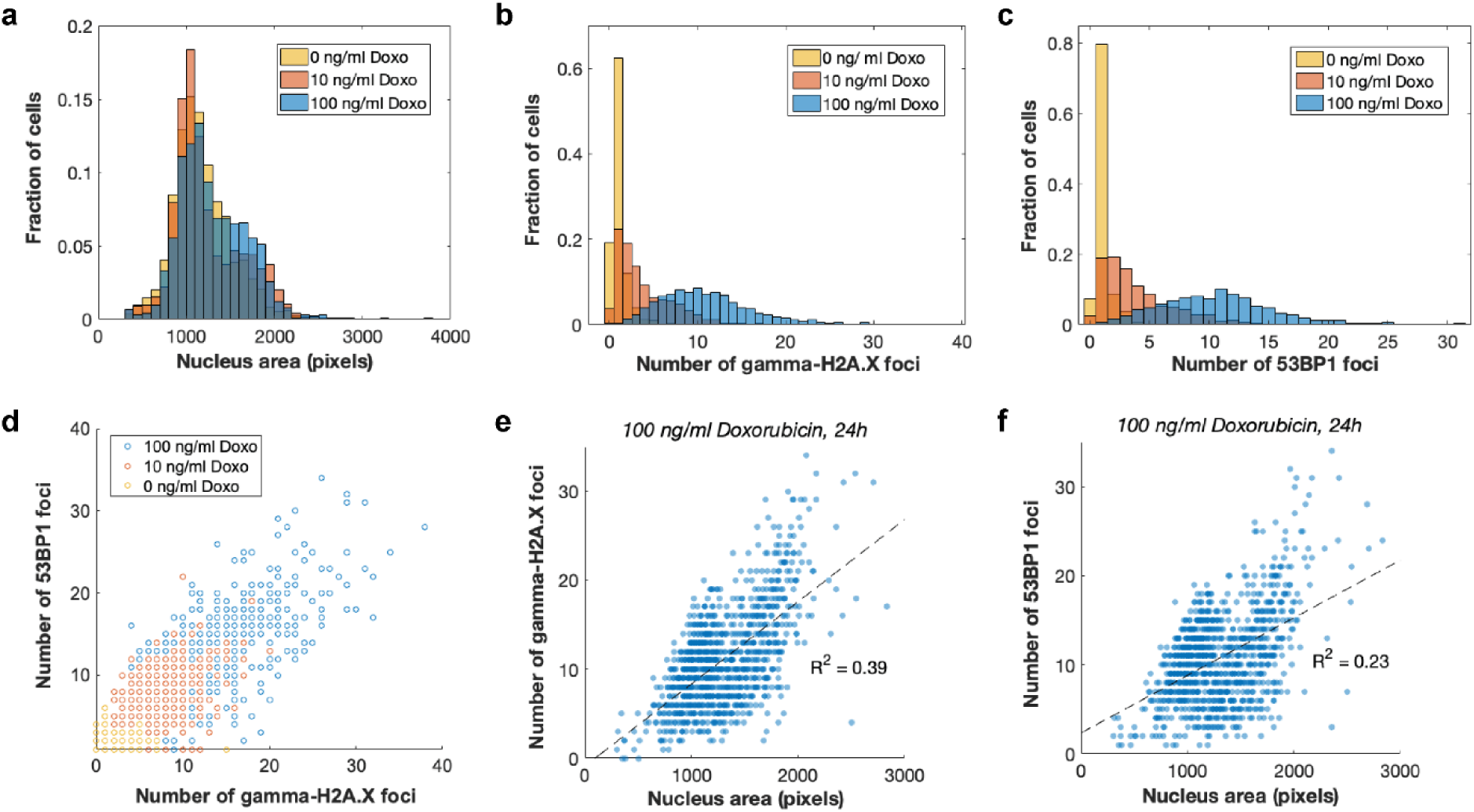
Correlation between the cell size and the number of DNA damage foci in RPE-1 cells. **(a-c)** Distribution of nuclear areas (a) and numbers of γ-H2AX (b) and 53BP1 (c) loci per nucleus in RPE-1 cells treated with DMSO, 10 ng/ml or 100 ng/ml Doxorubicin for 24 hours. **(d)** Correlation between the γ-H2AX and 53BP1 loci numbers in each RPE-1 nucleus. **(e-f)** Correlation between the nuclear area and the number of γ-H2AX (e) and 53BP1 (f) loci per nucleus in each RPE-1 cell nucleus after treatment with 100 ng/ml Doxorubicin for 24 hours. n = 1602 cells for DMSO, n = 1265 cells for 10 ng/ml Doxorubicin, and n = 1062 cells for 100 ng/ml Doxorubicin.

**Data S1. Protein and Peptide Slope values for size-sorted G1 cells**

**Data S2. Long-term Palbociclib time course**

**Data S3. Scaling behavior of protein complexes and their associated subunits**

**Data S4. mRNA concentrations in G1 HLF cells sorted into three size bins**

**Data S5. Ploidy-sorted Protein Slopes**

**Data S6. Experiment List and MaxQuant spectral search parameter file**

## References

1 Zatulovskiy, E. & Skotheim, J. M. On the Molecular Mechanisms Regulating Animal Cell Size Homeostasis. Trends Genet 36, 360–372, doi:10.1016/j.tig.2020.01.011 (2020).

2 Ginzberg, M. B., Kafri, R. & Kirschner, M. Cell biology. On being the right (cell) size. Science 348, 1245075, doi:10.1126/science.1245075 (2015).

3 Lloyd, A. C. The regulation of cell size. Cell 154, 1194–1205, doi:10.1016/j.cell.2013.08.053 (2013).

4 Lengefeld, J. et al. Cell size is a determinant of stem cell potential during aging. bioRxiv, 2020.2010.2027.355388, doi:10.1101/2020.10.27.355388 (2020).

5 Miettinen, T. P. & Bjorklund, M. Cellular Allometry of Mitochondrial Functionality Establishes the Optimal Cell Size. Dev Cell 39, 370–382, doi:10.1016/j.devcel.2016.09.004 (2016).

6 Neurohr, G. E. et al. Excessive Cell Growth Causes Cytoplasm Dilution And Contributes to Senescence. Cell 176, 1083–1097 e1018, doi:10.1016/j.cell.2019.01.018 (2019).

7 Padovan-Merhar, O. et al. Single mammalian cells compensate for differences in cellular volume and DNA copy number through independent global transcriptional mechanisms. Mol Cell 58, 339–352, doi:10.1016/j.molcel.2015.03.005 (2015).

8 Crissman, H. A. & Steinkamp, J. A. Rapid, simultaneous measurement of DNA, protein, and cell volume in single cells from large mammalian cell populations. J Cell Biol 59, 766–771, doi:10.1083/jcb.59.3.766 (1973).

9 Kempe, H., Schwabe, A., Cremazy, F., Verschure, P. J. & Bruggeman, F. J. The volumes and transcript counts of single cells reveal concentration homeostasis and capture biological noise. Mol Biol Cell 26, 797–804, doi:10.1091/mbc.E14-08-1296 (2015).

10 Lin, J. & Amir, A. Homeostasis of protein and mRNA concentrations in growing cells. Nat Commun 9, 4496, doi:10.1038/s41467-018-06714-z (2018).

11 Williamson, D. H. & Scopes, A. W. The distribution of nucleic acids and protein between different sized yeast cells. Exp Cell Res 24, 151–153, doi:10.1016/0014-4827(61)90258-0 (1961).

12 Zlotek-Zlotkiewicz, E., Monnier, S., Cappello, G., Le Berre, M. & Piel, M. Optical volume and mass measurements show that mammalian cells swell during mitosis. J Cell Biol 211, 765–774, doi:10.1083/jcb.201505056 (2015).

13 Son, S. et al. Resonant microchannel volume and mass measurements show that suspended cells swell during mitosis. J Cell Biol 211, 757–763, doi:10.1083/jcb.201505058 (2015).

14 Berenson, D. F., Zatulovskiy, E., Xie, S. & Skotheim, J. M. Constitutive expression of a fluorescent protein reports the size of live human cells. Mol Biol Cell 30, 2985–2995, doi:10.1091/mbc.E19-03-0171 (2019).

15 Chan, Y. H. & Marshall, W. F. Scaling properties of cell and organelle size. Organogenesis 6, 88–96, doi:10.4161/org.6.2.11464 (2010).

16 Marshall, W. F. Scaling of Subcellular Structures. Annu Rev Cell Dev Biol 36, 219–236, doi:10.1146/annurev-cellbio-020520-113246 (2020).

17 Cantwell, H. & Nurse, P. Unravelling nuclear size control. Curr Genet 65, 1281–1285, doi:10.1007/s00294-019-00999-3 (2019).

18 Zhurinsky, J. et al. A coordinated global control over cellular transcription. Curr Biol 20, 2010–2015, doi:10.1016/j.cub.2010.10.002 (2010).

19 Demidenko, Z. N. & Blagosklonny, M. V. Growth stimulation leads to cellular senescence when the cell cycle is blocked. Cell Cycle 7, 3355–3361, doi:10.4161/cc.7.21.6919 (2008).

20 Schmoller, K. M., Turner, J. J., Koivomagi, M. & Skotheim, J. M. Dilution of the cell cycle inhibitor Whi5 controls budding-yeast cell size. Nature 526, 268–272, doi:10.1038/nature14908 (2015).

21 Zatulovskiy, E., Zhang, S., Berenson, D. F., Topacio, B. R. & Skotheim, J. M. Cell growth dilutes the cell cycle inhibitor Rb to trigger cell division. Science 369, 466–471, doi:10.1126/science.aaz6213 (2020).

22 D’Ario, M. et al. Cell size controlled in plants using DNA content as an internal scale. Science 372, 1176–1181, doi:10.1126/science.abb4348 (2021).

23 Keifenheim, D. et al. Size-Dependent Expression of the Mitotic Activator Cdc25 Suggests a Mechanism of Size Control in Fission Yeast. Curr Biol 27, 1491–1497 e1494, doi:10.1016/j.cub.2017.04.016 (2017).

24 Chen, Y., Zhao, G., Zahumensky, J., Honey, S. & Futcher, B. Differential Scaling of Gene Expression with Cell Size May Explain Size Control in Budding Yeast. Mol Cell 78, 359–370 e356, doi:10.1016/j.molcel.2020.03.012 (2020).

25 Claude, K.-L., Bureik, D., Adarska, P., Singh, A. & Schmoller, K. M. Transcription coordinates histone amounts and genome content. bioRxiv, 2020.2008.2028.272492, doi:10.1101/2020.08.28.272492 (2020).

26 Swaffer, M. P. et al. Size-independent mRNA synthesis and chromatin-based partitioning mechanisms generate and maintain constant amounts of protein per cell. bioRxiv, 2020.2008.2028.272690, doi:10.1101/2020.08.28.272690 (2020).

27 Tzur, A., Moore, J. K., Jorgensen, P., Shapiro, H. M. & Kirschner, M. W. Optimizing optical flow cytometry for cell volume-based sorting and analysis. PLoS One 6, e16053, doi:10.1371/journal.pone.0016053 (2011).

28 Tan, C. et al. Cell size homeostasis is maintained by CDK4-dependent activation of p38 MAPK. Dev Cell 56, 1756–1769 e1757, doi:10.1016/j.devcel.2021.04.030 (2021).

29 Cheng, L. et al. Size-scaling promotes senescence-like changes in proteome and organelle content. bioRxiv, 2021.2008.2005.455193, doi:10.1101/2021.08.05.455193 (2021).

30 Liu, Y., Beyer, A. & Aebersold, R. On the Dependency of Cellular Protein Levels on mRNA Abundance. Cell 165, 535–550, doi:10.1016/j.cell.2016.03.014 (2016).

31 Zecha, J. et al. Peptide Level Turnover Measurements Enable the Study of Proteoform Dynamics. Mol Cell Proteomics 17, 974–992, doi:10.1074/mcp.RA118.000583 (2018).

32 Hernandez-Segura, A., Nehme, J. & Demaria, M. Hallmarks of Cellular Senescence. Trends Cell Biol 28, 436–453, doi:10.1016/j.tcb.2018.02.001 (2018).

33 Ruscetti, M. et al. NK cell-mediated cytotoxicity contributes to tumor control by a cytostatic drug combination. Science 362, 1416–1422, doi:10.1126/science.aas9090 (2018).

34 Demidenko, Z. N. et al. Rapamycin decelerates cellular senescence. Cell Cycle 8, 1888–1895, doi:10.4161/cc.8.12.8606 (2009).

35 Sharpless, N. E. & Sherr, C. J. Forging a signature of in vivo senescence. Nat Rev Cancer 15, 397–408, doi:10.1038/nrc3960 (2015).

36 Sage, J. et al. Targeted disruption of the three Rb-related genes leads to loss of G(1) control and immortalization. Genes Dev 14, 3037–3050, doi:10.1101/gad.843200 (2000).

37 Dannenberg, J. H., van Rossum, A., Schuijff, L. & te Riele, H. Ablation of the retinoblastoma gene family deregulates G(1) control causing immortalization and increased cell turnover under growth-restricting conditions. Genes Dev 14, 3051–3064, doi:10.1101/gad.847700 (2000).

38 Mu, L. et al. Mass measurements during lymphocytic leukemia cell polyploidization decouple cell cycle- and cell size-dependent growth. Proc Natl Acad Sci U S A 117, 15659–15665, doi:10.1073/pnas.1922197117 (2020).

39 Lee, H. O., Davidson, J. M. & Duronio, R. J. Endoreplication: polyploidy with purpose. Genes Dev 23, 2461–2477, doi:10.1101/gad.1829209 (2009).

40 Kumari, R. & Jat, P. Mechanisms of Cellular Senescence: Cell Cycle Arrest and Senescence Associated Secretory Phenotype. Front Cell Dev Biol 9, 645593, doi:10.3389/fcell.2021.645593 (2021).

41 Sakaue-Sawano, A. et al. Visualizing spatiotemporal dynamics of multicellular cell-cycle progression. Cell 132, 487–498, doi:10.1016/j.cell.2007.12.033 (2008).

42 Deng, J., Erdjument-Bromage, H. & Neubert, T. A. Quantitative Comparison of Proteomes Using SILAC. Curr Protoc Protein Sci 95, e74, doi:10.1002/cpps.74 (2019).

43 Zecha, J. et al. TMT Labeling for the Masses: A Robust and Cost-efficient, In-solution Labeling Approach. Mol Cell Proteomics 18, 1468–1478, doi:10.1074/mcp.TIR119.001385 (2019).

44 Cox, J. et al. Andromeda: a peptide search engine integrated into the MaxQuant environment. J Proteome Res 10, 1794–1805, doi:10.1021/pr101065j (2011).

45 Cox, J. & Mann, M. MaxQuant enables high peptide identification rates, individualized p.p.b.- range mass accuracies and proteome-wide protein quantification. Nat Biotechnol 26, 1367–1372, doi:10.1038/nbt.1511 (2008).

46 Elias, J. E. & Gygi, S. P. Target-decoy search strategy for increased confidence in large-scale protein identifications by mass spectrometry. Nat Methods 4, 207–214, doi:10.1038/nmeth1019 (2007).

47 UniProt, C. UniProt: a worldwide hub of protein knowledge. Nucleic Acids Res 47, D506–D515, doi:10.1093/nar/gky1049 (2019).

48 Cox, J. & Mann, M. 1D and 2D annotation enrichment: a statistical method integrating quantitative proteomics with complementary high-throughput data. BMC Bioinformatics 13 **Suppl 16**, S12, doi:10.1186/1471-2105-13-S16-S12 (2012).

49 Frenkel-Morgenstern, M. et al. Genes adopt non-optimal codon usage to generate cell cycle-dependent oscillations in protein levels. Mol Syst Biol 8, 572, doi:10.1038/msb.2012.3 (2012).

50 Frankish, A. et al. GENCODE reference annotation for the human and mouse genomes. Nucleic Acids Res 47, D766–D773, doi:10.1093/nar/gky955 (2019).

51 Yates, A. D. et al. Ensembl 2020. Nucleic Acids Res 48, D682–D688, doi:10.1093/nar/gkz966 (2020).

52 Dobin, A. et al. STAR: ultrafast universal RNA-seq aligner. Bioinformatics 29, 15–21, doi:10.1093/bioinformatics/bts635 (2013).

53 Langmead, B., Trapnell, C., Pop, M. & Salzberg, S. L. Ultrafast and memory-efficient alignment of short DNA sequences to the human genome. Genome Biol 10, R25, doi:10.1186/gb-2009-10-3-r25 (2009).

54 Roberts, A. & Pachter, L. Streaming fragment assignment for real-time analysis of sequencing experiments. Nat Methods 10, 71–73, doi:10.1038/nmeth.2251 (2013).

55 Schwarz, C. et al. A Precise Cdk Activity Threshold Determines Passage through the Restriction Point. Mol Cell 69, 253–264 e255, doi:10.1016/j.molcel.2017.12.017 (2018).

